# Identification of Relevant Genetic Alterations in Cancer using Topological Data Analysis

**DOI:** 10.1101/2020.01.30.922310

**Authors:** Raúl Rabadán, Yamina Mohamedi, Udi Rubin, Tim Chu, Oliver Elliott, Luis Arnés, Santiago Cal, Álvaro J. Obaya, Arnold J. Levine, Pablo G. Cámara

## Abstract

Large-scale cancer genomic studies enable the systematic identification of mutations that lead to the genesis and progression of tumors, uncovering the underlying molecular mechanisms and potential therapies. While some such mutations are recurrently found in many tumors, many others exist solely within a few samples, precluding detection by conventional recurrence-based statistical approaches. Integrated analysis of somatic mutations and RNA expression data across 12 tumor types reveals that mutations of cancer genes are usually accompanied by substantial changes in expression. We use topological data analysis to leverage this observation and uncover 38 elusive candidate cancer-associated genes, including inactivating mutations of the metalloproteinase ADAMTS12 in lung adenocarcinoma. We show that *ADAMTS12*^−/−^ mice have a five-fold increase in the susceptibility to develop lung tumors, confirming the role of *ADAMTS12* as a tumor suppressor gene. Our results demonstrate that data integration through topological techniques can increase our ability to identify previously unreported cancer-related alterations.

## Introduction

A critical foundation for targeted cancer therapies is the identification of molecular mechanisms that are necessary for tumor development and maintenance. Large-scale crosssectional cancer molecular studies, such as The Cancer Genome Atlas (TCGA) and the International Cancer Genome Consortium, enable this identification by systematically compiling genetic alterations across many tumors^1,2^. Tumors that present recurrently altered genes or pathways are suspected to be driven by common molecular mechanisms. By leveraging computational approaches that seek signatures of positive selection^3^, these studies have produced extensive catalogues of frequently-mutated, cancer-associated genes^4^. These studies have also revealed that most cancer mutations occur at low frequencies (< 10% of samples), including potentially actionable therapeutic targets^4^.

The identification of low-prevalence cancer-associated mutations using recurrence-based methods is challenging because of the large number of samples needed to achieve statistical power and the inherent complexity in modeling the background mutation rates. The rate of neutral mutations within a cancer type can dramatically differ among patients, genomic regions, or mutation types^3^, limiting the power of recurrence-based methods. It is estimated that for the current size of ongoing cross-sectional studies (typically consisting of less than 1,000 patients) only cancer-associated mutations that occur at intermediate or high frequencies (> 15%) are fully accessible to recurrence-based methods^4^. This is consistent with the observation that patients in these studies often completely lack mutations in known cancer-associated genes^5^. These results highlight the need of recurrence-based methods that can model rare events^6,7^ or, alternatively, methods for the identification of cancer-associated genes that are not based on recurrence.

An approach to the identification of cancer-associated genes that is not based on modeling the mutation rate is the integration of other types of data from the tumor^8,9^. If a mutation is accompanied by consistent changes in copy number, gene expression, and/or methylation, it is possible to leverage these changes to relate the mutation event to cancer progression. Several studies have utilized changes in the copy number, expression, or methylation of the mutated gene (*cis*-effects) to identify novel cancer-associated mutations^10–12^. However, the identification of cancer-associated mutations based on changes in genes other than the mutated gene (*trans*-effects) is more challenging and usually requires providing information about known gene-gene relationships to reduce the number of false positives. DriverNet^13^, OncoIMPACT^14^, CaMoDi^15^, and Xseq^16^ utilize *trans*-effects in gene expression to identify cancer-associated mutations. To limit the dimensionality of the expression space, these methods use expression modules (sets of co-expressed genes, functionally related genes, or gene networks). However, an approach that fully takes into account the complexity of the expression space is currently lacking.

Current approaches for the identification of cancer-associated mutations using expression data are sensitive to several confounding effects. Genomic regions with open chromatin can be more easily accessed by DNA repair enzymes, leading to anti-correlations between gene expression levels and mutation rates^3^. Furthermore, tumors with different expression signatures, such as genomically unstable tumors, can have different mutation rates^17,18^. These effects lead to spurious correlations between mutations and expression signatures and are a source of false positives for current algorithms.

To address these problems, we have devised an approach to identify cancer-associated mutated genes using expression data from multiple tumors. Our approach makes use of topological data analysis^19,20^ (TDA) to reconstruct the structure of the expression space, and takes into account the above spurious effects when assessing the significance of a mutated gene. Its application to mutation and expression data of 4,476 patients from 12 tumor types leads to the identification of 95 mutated cancer genes, out of which 38 are previously unreported low-prevalence genes (average prevalence within the same tumor cohort = 5%). We hence propose a complementary approach to recurrence-based methods, enabling the identification of elusive, but potentially clinically-relevant, mutated cancer genes.

## Results

### Topological reconstruction of the expression space of low grade gliomas

The expression profile of a tumor can be mathematically described as a point in a highdimensional expression space, where each dimension represents the mRNA level of a gene and the dimensionality of the space is given by the number of expressed genes. Points that lie close to each other in this space correspond to tumors with similar expression profiles. The set of all possible tumors of a cancer type spans a sub-space of the expression space. Measuring the expression profiles of individual tumors in a cross-sectional study is equivalent to sampling a finite set of points from this sub-space.

We considered 513 primary low grade glioma (LGG) tumors from TCGA for which both RNA-seq and whole-exome DNA-seq data were available^21^. To infer the structure of the expression space of LGG from this RNA-seq data, we used a topological approach^19,20^. Topology is the mathematical field that studies how different parts of a space are connected to each other. TDA generalizes some of the notions of topology to sets of points and pairwise distances. Thus, TDA aims to infer and summarize the topological structure of a space given only a finite sample of points. TDA has been recently used to study viral re-assortment^22^, human recombination^23,24^, cell differentiation^25^, breast cancer^26^, and other complex genetic diseases^27^.

We used the TDA algorithm Mapper^28^ to build a low-dimensional representation of the expression space of LGG using the expression data of the TCGA cohort (Fig. 1a). Mapper generates a network representation of the expression space, in which each node corresponds to a set of tumors with similar expression profile. A given tumor can appear in more than one node, and if two nodes have one or more tumors in common they are connected by an edge. Contrary to other methods for dimensionality reduction, such as principal component analysis and multi-dimensional scaling^29^, the topological representations produced by Mapper preserve local relationships of the high-dimensional expression space. Any two tumors close to each other in the topological representation (as measured by the number of edges contained in the shortest path that connects the two tumors) are ensured to be close to each other in the original high-dimensional expression space. We used Pearson’s correlation as a measure of similarity between the expression profiles of individual tumors. The topological representation of the LGG expression space consisted of three regions (Fig. 1a), consistent with the expression sub-types found in clustering analyses^21^. These regions, however, were bridged by thin structures in the topological representation, indicating that some tumors have an expression profile characteristic of multiple expression sub-types (Fig. 1a).

**Figure 1.**
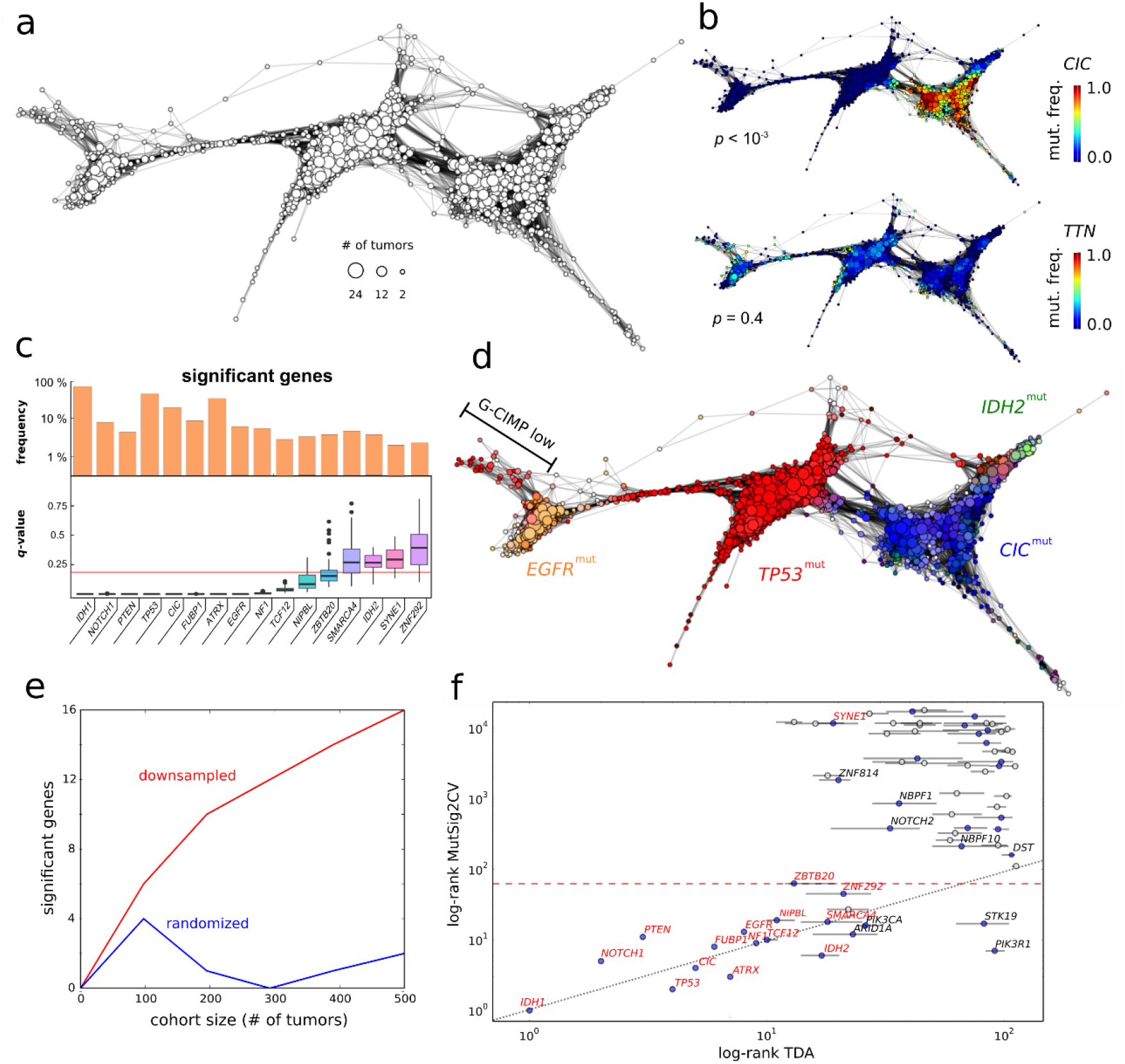
Identification of mutated cancer genes in LGG using an integrative topological approach. (**a**) Topological representation of the expression space of LGG based on the expression data of 513 tumors. Each node represents a set of tumors with similar global expression patterns. The size of each node represents the number of tumors in the set. Edges connect nodes that share at least one tumor. Three large expression groups are clearly visible in the representation. (**b**) The localization of each mutated gene in the topological representation is assessed statistically. Mutated genes significantly localized in the expression space are candidate drivers of tumor progression. The topological representation of the expression space of LGG labeled by the frequency of somatic mutations of the *CIC* (top) and *TTN* (bottom) genes is displayed as an example. *CIC* mutations are significantly localized in the expression space of LGG, consistently with being a driver of tumor progression. (**c**) List of significantly (*q*-value < 0.15) localized mutated genes in the reconstructed expression space of LGG. The prevalence of mutations in the cohort and the distribution of the statistical significance across the parameter space of the topological representation are also displayed. Box-plot elements: center line, median; box limits, upper and lower quartiles; whiskers, 1.5x interquartile range; points, outliers. (**d**) The topological representation of the expression space of LGG is labeled according to the prevalence of some of the significantly localized mutations. The three large expression groups in the topological representation are identified with oligodendrogliomas (enriched for *CIC* and *IDH2* mutations), *IDH1*-mutant astrocytomas (enriched for *TP53* mutations), and *IDH1*-wild-type astrocytomas (enriched for *EGFR* mutations). *IDH1*-mutant astrocytomas with a low G-C island methylation phenotype (G-CIMP low) form a flare of *IDH1*-wild-type astrocytomas. (**e**) Number of significant mutated genes as a function of the cohort size, for the original (red) and a randomized (blue) version of the LGG cohort. Our integrative topological approach produces significant results for tumor cohorts above ~100 patients. (**f**) Comparison of the results of the integrative topological approach with those of MutSig2CV on the same cohort. Represented is the rank of each gene according to their significance in the topological (horizontal axis) and MutSig2CV (vertical axis) analyses in logarithmic scale. Genes that are significant (*q*-value < 0.15) according to our topological approach are marked in red. Genes below the red dashed line are significant (*q*-value < 0.15) according to the MutSig2CV analysis. Horizontal bars indicate the 16% and 84% percentiles of the ranks across the parameter space of the topological representation. Besides being radically different approaches, the results of our topological approach and MutSig2CV display a large degree of consistency, with most cancer-associated mutated genes sitting across the diagonal.

### Identification of cancer-associated mutated genes in LGG

We hypothesized that if a mutated gene appears localized in the expression space, it is associated with consistent global expression patterns across a subset of tumors, and is therefore a candidate driver of tumor progression (Fig. 1b). On the other hand, if mutations of a gene are clonally expanded as a result of being in the same genome as a positively-selected mutation, but are not cancer-related, they will appear randomly scattered in the expression sub-space (Fig. 1b).

To test this hypothesis, we implemented a computational approach that assesses the localization of non-synonymous somatic mutations in the expression space of tumors (Methods). To control for the presence of spurious correlations between the mutation rate and the tumor expression profile, we assessed the localization of the mutational tumor burden (defined as the total number of somatic mutations in each tumor) in the reconstructed expression space (Supplementary Fig. 1a, Methods). Based on this analysis, we sub-sampled mutations in two hyper-mutated tumors (n_mut_ > 10^2.5^) that were present in the LGG cohort. In addition, we assessed the similarity between the expression and the mutation profile of each individual gene in the reconstructed expression space (Methods). After correcting for these spurious correlations, 16 mutated genes were significantly localized in the reconstructed expression space of LGG (Fig. 1c, *q*-value < 0.15, Benjamini-Hochberg procedure). These included well-known high-prevalence (> 15%) driver genes, like *IDH1*, *TP53*, *ATRX*, and *CIC*, in addition to several low-prevalence mutated genes, like *NIPBL* (mutated in 4% of the tumors) and *ZNF292* (mutated in 3% of the tumors), which have been recently reported in a larger cohort of gliomas^21^. In total, 15 out of the 16 significant mutated genes were previously reported^21,30^, with *SYNE1* (mutated in 2% of the tumors) the only new candidate. We did not observe a significant correlation between the significance and prevalence of statistically significant genes (Pearson’s correlation coefficient between prevalence and *q*-value, *r* = −0.34, *p*-value = 0.19). In particular, some of the most significant genes according to our approach, like *FUBP1*, *NOTCH1*, *PTEN*, *EGFR*, and *NF1*, were mutated in less than 10% of the patients within that tumor type (Fig. 1c), indicating that mutations in these genes are strongly associated with global changes in expression. These results were stable across the parameter space of the Mapper algorithm (Fig. 1c, Supplementary Fig. 1b, Methods).

The location of significant genes in the reconstructed expression space of LGG was consistent with the known molecular subtypes of adult diffuse gliomas^21^ (Fig. 1d, Supplementary Fig. 2). Of particular note, *IDH2*-mutant tumors were localized within the expression space of oligodendrogliomas, indicating a distinct expression profile from that of *IDH1*-mutant oligodendrogliomas (Fig. 1d, Supplementary Fig. 2a). This observation is consistent with a recent study based on genomic variations^31^.

Neuronal marker expression has been reported in malignant (grade III/IV) gliomas other than classical anaplastic gangliogliomas^32,33^. In our cohort, tumors expressing canonical neuronal markers like neurofilament (*NEFL*, *NEFM*, and *NEFH*) and synaptophysin (*SYP*) were significantly localized within the expression space of oligodendrogliomas (*q*-value < 0.015, Supplementary Fig. 3). These tumors harbored frequent deletions of the chromosome arm 19q, in addition to molecular alterations characteristic of astrocytic gliomas, such as *TP53* and *ATRX* mutations (Fisher’s exact test *p*-value < 0.01, Supplementary Fig. 3). Although the average estimated tumor purity^34^ in this group was significantly lower than for the rest of the oligodendroglioma expression group (Mann-Whitney U-test *p*-value = 0.001, average estimated tumor purities = 92% and 96%, respectively), the estimated tumor content was in many cases (*n* = 7) above 98%, suggesting that the expression of neuronal markers is not due to a poor tumor purity.

### Computational benchmarking

To assess the number cancer-associated genes identified by our approach as a function of the size of the cohort, we repeated the same analysis in smaller cohorts generated by randomly sampling patients from the original LGG cohort (Fig 1e). We also assessed the number of false positives by generating randomized datasets, where we permuted the labels of the patients in the expression data. We observed that our approach requires a minimum cohort size of approximately 100 tumors. For larger cohorts, the expected number of false positives was between 1 and 2 (Fig. 1e).

Next, we sought to compare our results against current algorithms for the identification of cancer-associated genes using expression data. To that end, we analyzed the same LGG cohort using the recently published algorithm Xseq^16^ (Methods). Xseq implements a hierarchical Bayes statistical model to quantify the impact of somatic mutations on expression profiles using a pre-computed ‘influence graph’ that encodes whether two genes are known to be functionally related. The analysis of the LGG cohort with Xseq led to only 2 significant genes (posterior probability, *P(D)* > 0.80), of which only one (*PTEN*) has been previously reported in LGG. These results reveal the high sensitivity of our topological approach compared to state-of-the-art algorithms.

In addition to Xseq, we compared the results of our integrative topological approach to those produced by MutSig2CV on the same cohort (Fig. 1f). MutSig2CV models the neutral background mutation rate, taking into account genomic variations due to differences in expression level and replication time^3^. We observed a significant overlap between the results of MutSig2CV and those of our approach, with 15 out of 23 mutated genes that were significant (*q*-value < 0.15) according to MutSig2CV, being also significant according to our approach (65% overlap, Fisher’s exact test *p*-value = 10^−42^). Some of the most significant cancer genes identified by MutSig2CV based on recurrence, such as *PIK3R1* (mutated in 4% of the tumors), were not selected by our expression-based approach, highlighting the independence of recurrence- and expression-based approaches. Combining the results of MutSig2CV with those of our integrative topological approach (Fig. 1f) singled out new low-prevalence mutated genes, such as *NOTCH2*, as potential drivers of tumor progression in LGG.

Seeking a more systematic comparison with existing methods, we performed a similar study to that of Bertrand *et al*.^35^ across multiple tumor types (Methods). We estimated the precision, recall, and F1 score of our integrative topological approach, Xseq, MutSig2CV, OncodriveFML^36^, and 20/20+^37^ based on the overlap of their top 15 predictions with a gold-standard list of cancer-associated genes^35^. In addition to the LGG cohort, we analyzed two cohorts of 208 colorectal adenocarcinoma (COAD) and 930 breast invasive carcinoma (BRCA) tumors from TCGA, respectively. In each of the three cohorts, the precision, recall, and F1 score of our integrative topological approach were the highest or second highest among the 5 algorithms (Supplementary Table 1), highlighting its utility for the identification of mutated cancer-associated genes.

### Identification of cancer-associated genes across 12 tumor types

Based on the above results, we decided to extend our analysis to other tumor types. We considered 12 tumor types from TCGA for which there were sufficient samples (*n* > 140) with RNA-seq and whole-exome data available (Table 1). The complete results of our analysis can be accessed through an online database (Methods). In total, our approach identified 95 mutated cancer genes (*q*-value < 0.15), out of which 16 genes were significant in two or more tumor types (Fig. 2a, Supplementary Figs. 4 – 15, and Supplementary Table 2). Some of the most common genes were *TP53*, *KRAS*, *HRAS*, *PIK3CA*, *ATRX*, *EGFR*, and *NF1*. The number of significant genes in each tumor type was correlated with the size of the cohort (Fig. 2b, Spearman’s correlation coefficient *r* = 0.67, *p*-value = 0.02), consistently with the results of the computational benchmarking. We observed a large degree of consistency between the list of significant genes and curated databases of cancer genes. Specifically, 61% of the significant genes in our analysis were present in the Cancer Gene Census^38^ or OncoKB^39^ databases (Fig. 2c, Fisher’s exact test *p*-value < 10^−50^ for each database). Overall, 75% of the patients carried a mutation in a significant gene, out of which 24% carried mutations in actionable genes with approved drugs^35^ (Supplementary Table 3).

**Figure 2.**
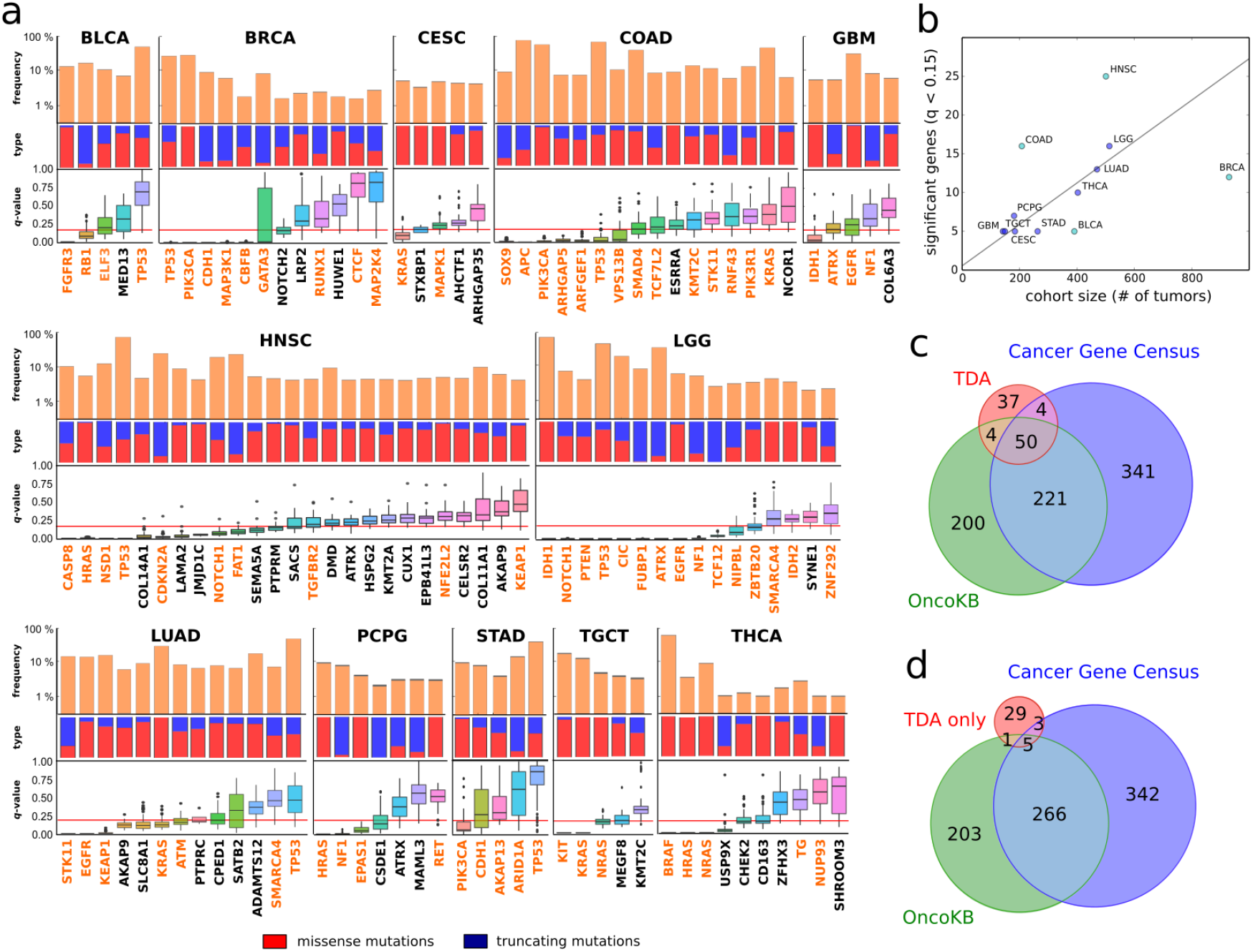
Identification of mutated cancer genes across 12 tumor types using an integrative topological approach. (**a**) Significant genes (*q*-value < 0.15) in the integrative topological analysis of the 12 tumor types considered in Table 1. From top to bottom, the frequency of non-synonymous mutations, the fraction of missense versus truncating mutations, and the distribution of *q*-values across the parameter space is shown for each gene. Genes that are also significant (*q*-value < 0.15) based on MutSig2CV are shown in orange. Box-plot elements: center line, median; box limits, upper and lower quartiles; whiskers, 1.5x interquartile range; points, outliers. (**b**) Plot of the number of significant genes (*q*-value < 0.15) against the number of tumors in the cohort, for each tumor type. A linear fit is shown, where outliers (marked in cyan) where not taken into account in the fit (Pearson’s *r* = 0.94, *p*-value = 5·10^−4^). (c) Venn diagram showing the overlap between significant genes (TDA) and the curated databases of cancer genes OncoKB (Fisher’s exact test *p*-value < 10^−50^) and the Cancer Gene Census (Fisher’s exact test *p*-value < 10^−50^). (**d**) Venn diagram showing the overlap between genes that are significant in the integrative topological analysis but not in the MutSig2CV analysis (TDA only), and the curated databases of cancer genes OncoKB (Fisher’s exact test *p*-value = 3·10^−4^) and the Cancer Gene Census (Fisher’s exact test *p*-value = 2·10^−5^).

**Table 1.** Number of patients in each of the cohorts analyzed using topological (TDA) and recurrence-based (MutSig2CV) approaches.

| Cancer Type | Cohort | Samples<br>(TDA) | Samples<br>(MutSig2CV) |
| --- | --- | --- | --- |
| Bladder urothelial carcinoma | BLCA | 391 | 395 |
| Breast invasive carcinoma | BRCA | 930 | 978 |
| Cervical and endocervical cancers | CESC | 184 | 194 |
| Colon adenocarcinoma | COAD | 208 | 367 |
| Glioblastoma multiforme | GBM | 142 | 283 |
| Head and neck squamous cell carcinoma | HNSC | 501 | 511 |
| Brain lower grade glioma | LGG | 513 | 516 |
| Lung adenocarcinoma | LUAD | 470 | 533 |
| Pheochromocytoma and paraganglioma | PCPG | 181 | 179 |
| Stomach adenocarcinoma | STAD | 263 | 393 |
| Testicular germ cell tumors | TGCT | 149 | 147 |
| Thyroid carcinoma | THCA | 403 | 496 |
| **Total:** |  | **4,335** | **4,992** |

The results were largely consistent with those of MutSig2CV on the same TCGA cohorts (Fig. 2a, Supplementary Fig. 16, and Table 1), adding further support to some of the cancer genes identified in our analysis. Out of the 95 significant genes in the integrative topological analysis, 38 genes were not significant according to MutSig2CV (Fig. 2a, *q*-value < 0.15). However, these putative elusive cancer genes did often displayed a tendency towards significance in the MutSig2CV analysis, likely reflecting a limitation of the cohort size (Supplementary Fig. 16). They also had a significant overlap with the Cancer Gene Census (8 out of 38 genes, Fisher’s exact test *p*-value = 2·10^−5^, Fig. 2d) and OncoKB (6 out of 38 genes, Fisher’s exact test *p*-value = 3·10^−4^, Fig. 2d) databases, as well as with genes involved in developmental processes (27 out of 38 genes, g:SCS *q*-value = 10^−3^). Elusive genes included *NOTCH2* mutations in breast invasive carcinoma (mutated in 2% of the tumors), which have been recently reported by manual inspection^40^; inactivating mutations of the tumor-suppressor genes *KMT2A^41^* (also known as *MLL1*) and *CUX1^42^* in head and neck squamous cell carcinoma (each present in 1% of the tumors); inactivating mutations of the tumor-suppressor gene *ADAMTS12^43^* in lung adenocarcinoma (present in 4% of the tumors); mutations in the kinase domain of *CHEK2* in thyroid carcinoma (present in 1% of the tumors), which have been associated with increased susceptibility to this cancer type^44^; inactivating mutations of the putative tumor-suppressor gene *USP9X* in thyroid carcinoma (present in 1% of the tumors), which codes for a deubiquitinase regulating the TGFβ^45^ and hippo signaling pathways^46^; and inactivating mutations of *ATRX* in pheochromocytoma and paraganglioma^47^ (present in 2% of the tumors). These genes, except *CHEK2*, encode long proteins (>1,500 amino-acids) and are expected to contain numerous passenger mutations, complicating the identification of low-prevalence cancer-associated mutations using recurrence-based methods.

Additionally, the combination of the results of our analysis with those of MutSig2CV allowed us to prioritize the study of mutated genes in colon adenocarcinoma, where the number of significant genes according to MutSig2CV is too large (*n* = 1,698 genes, *q*-value < 0.15) (Supplementary Fig. 16). In particular, our analysis highlighted *ARHGAP5* and *ARFGEF1* as previously unreported putative driver genes of tumor progression in this cancer type.

### Truncating mutations of *ADAMTS12* in lung adenocarcinoma (LUAD) are associated to poor survival in humans and increased tumor susceptibility in mice

Using TCGA survival data we found that, among the previously unreported cancer-associated genes, inactivating mutations of ADAMTS12 were associated with poor survival (Fig. 3a). ADAMTS12 is a metalloproteinase with thrombospondin motif that can block the activation of the Ras-MAPK signaling pathway^43^. Immunodeficient mice injected with A549 lung adenocarcinoma cells overexpressing ADAMTS12 had a deficiency of tumor growth in comparison with tumors formed from parental A549 cells^43^. The *ADAMTS12* gene is in chromosomal arm 5p, which is entirely amplified in over 60% of lung adenocarcinoma tumors^48^. It has been suggested that the *TERT* gene, coding for the telomerase catalytic subunit, may be the target of this amplification^48^. Consistent with the suggested anti-tumorgenic properties of ADAMTS12, we observed that LUAD patients with chromosome 5p amplification and unaltered *ADAMTS12* gene have better overall survival than those without chromosome 5p amplification (Fig. 3a, median overall survival 4.2 years versus 3.4 years respectively, Kaplan-Meier *p*-value = 0.05). To the contrary, patients with chromosome 5p amplification and truncating mutations in *ADAMTS12* have a reduced survival with respect to patients that harbor the amplification without mutations in *ADAMTS12* (Fig. 3a, median overall survival 2.4 years, Kaplan-Meier *p*-value = 0.015). Additionally, truncating mutations in *ADAMTS12* tend to co-occur with chromosome 5p amplification (Fig. 3a, one-tailed Fisher’s exact test *p*-value = 2·10^−3^).

**Figure 3.**
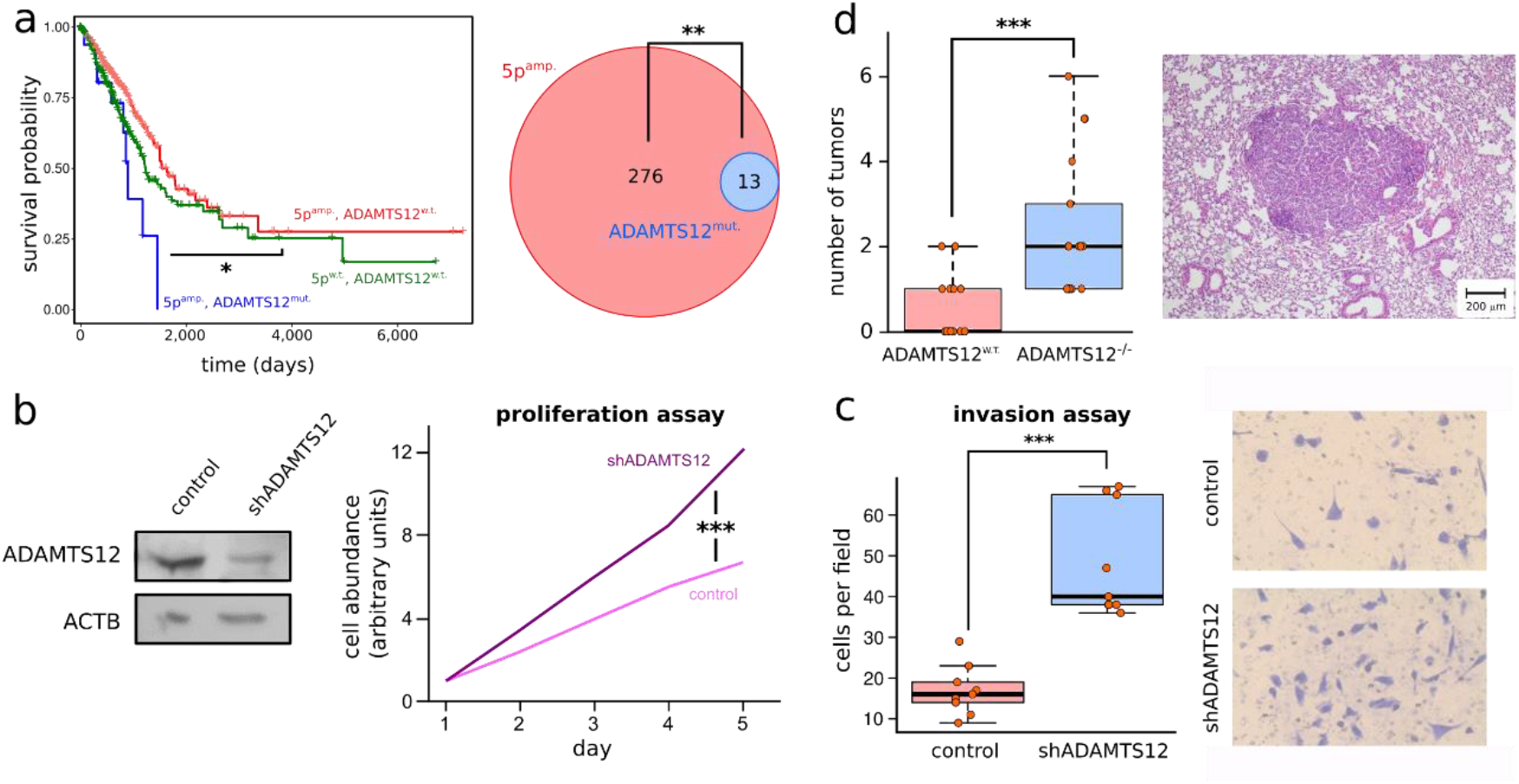
Truncating mutations of *ADAMTS12* are associated to increased tumor susceptibility and poor survival in LUAD. (**a**) Left: Kaplan-Meier survival curves for the LUAD cohort, where patients have been stratified according to whether their tumors have chromosome 5p amplification and absence of truncating mutations in *ADAMTS12* (red), chromosome 5p amplification and presence of truncating mutations in *ADAMTS12* (blue), and absence of both chromosome 5p amplification and truncating mutations in *ADAMTS12* (green). Kaplan-Meier *p*-values of blue and green survival curves with respect to red survival curve are 0.015 and 0.05 respectively. Right: Venn diagram showing the overlap between tumors with chromosome 5p amplification (red) and tumors with truncating mutations in *ADAMTS12* (blue) (one-tailed Fisher’s exact test *p*-value = 2·10^−3^). (**b**) Left: Western-blot showing the expression of ADAMTS12 and β-Actin in control and shADAMTS12 LL/C-luc-M38 cells. Right: In-vitro proliferation assay of control and shADAMTS12 cells. (**c**) Left: Number of control and shADAMTS12 cells with invasive potential in an in-vitro invasion assay. Right: Representative images of fields used for quantification after invasion assay. (**d**) Left: Number of lung tumors observed in control (*n* = 17) and ADAMTS12-deficient (*n* = 17) mice treated with urethane after 20 weeks of treatment. ADAMTS12-deficient mice show a 5-fold enrichment in the number of lung tumors as compared to control mice. Right: Hematoxylin-eosin stained tissue section of ADAMTS12-deficient mice treated with urethane displaying a lung adenocarcinoma tumor. Box-plot elements: center line, median; box limits, upper and lower quartiles; whiskers, 1.5x interquartile range; points, outliers. Source data are provided as a Source Data file.

To validate ADAMTS12 inactivation as a driver of progression in lung carcinoma, we investigated the effect of silencing ADAMTS12 in the lung carcinoma cell line LL/2-luc-M38 using a shRNA plasmid. In vitro proliferation and invasion assays revealed a significant increase in the proliferative and invasive potential of the cells that were transfected with the shRNA plasmid compared to the control cells (Figs. 3b, c, Mann-Whitney U-test *p*-value < 10^−3^ in both assays).

In addition to these in vitro studies, we assessed the effect of ADAMTS12 inactivation in vivo. To that end, we generated ADAMTS12^−/−^ mice as previously described^49^ and treated ADAMTS12 knockout and control mice with urethane (ethyl carbamate), a carcinogen that typically induces lung adenomas after several months of treatment^50,51^. After 20 weeks of treatment, ADAMTS12 knockout mice showed a 5-fold enrichment on the number of lung tumors compared to control mice (Fig. 3d, Mann-Whitney U-test *p*-value = 3·10^−5^). The enrichment in the number of tumors was still significant after disaggregating tumors by their size (Supplementary Fig. 17). We did not find a significant difference between the observed tumor size in control and ADAMTS12 knockout mice. Immunohistochemistry staining of tumor sections from these mice revealed some level of expression of ADAMTS12 in the region surrounding the adenoma, but not in the highly-proliferative Ki-67^+^ cells (Supplementary Fig. 18). A similar pattern of ADAMTS12 expression has been observed in human colon adenocarcinoma^52^. Taken together our results suggest ADAMTS12 has a tumor suppressor role in lung cancer, consistently with the results of our computational analysis.

## Discussion

To identify which somatic mutations are relevant to the progression of tumors, most genomic analyses focus on the recurrence of mutations and define candidate cancer-associated genes as those mutated at a higher frequency than expected under a modeled local neutral mutation rate. This definition has proven to be particularly powerful for commonly mutated genes. However, it is limiting for low prevalence mutations or tumors with a higher mutation burden. Here, we have adopted an alternative definition for candidate cancer-associated gene based on the assumption that mutations in these genes are accompanied by consistent global expression patterns in the tumor. Remarkably, these two fundamentally different definitions are in practice highly consistent with each other, as we find that most mutations occurring at a high frequency compared to the local neutral mutation rate are associated with consistent global mRNA expression patterns in the tumor. As expected, there are numerous exceptions to this rule and utilizing our approach we are able to identify multiple candidate cancer genes that remained elusive to other methods. One example of such elusive cancer-associated mutations are truncating mutations of the PEST domain of *NOTCH2* occurring in breast invasive carcinoma^40^. These rare events are easily masked by the large number of passenger mutations that this long gene accumulates. However, we find these alterations are consistently accompanied by global changes in the expression profile of the tumor. Although they affect a small fraction of all breast cancer patients, the availability of pharmacological inhibitors of the Notch signaling pathway makes them a promising therapeutic target for the treatment of these patients^53^. Among the less studied, elusive candidate cancer-associated mutations identified with our approach, we have studied the inactivating mutations of *ADAMTS12* occurring in lung adenocarcinoma. We have provided evidence of the tumor suppressor role of *ADAMTS12* in this cancer type both in vitro and in vivo. Specifically, our experiments reveal that lung carcinoma LL/2-luc-M38 cells display a higher proliferative and invasive potential in vitro when transfected with an ADAMTS12 shRNA. Additionally, we have shown that mice treated with urethane have a several fold increase in the susceptibility to develop lung adenomas when *ADAMTS12* is knocked out. These results are consistent with the observation that patients of lung adenocarcinoma with tumors harboring truncating mutations of *ADAMTS12* have poor survival. Our work demonstrates that the combination of recurrence-based methods with integrative approaches as we describe here can be a valuable tool to systematically identify potentially actionable, low-prevalence mutations that escape standard methods of detection.

## Methods

### Sample collection and preprocessing

We collected gene expression levels and somatic mutation data of 12 tumor types from the TCGA repository (http://cancergenome.nih.gov/) (Table 1 and Supplementary Table 4). We only considered patients for which both types of data were available. RNA-seq expression levels were retrieved in RSEM format^54^ and estimated relative abundances (*x*) were transformed according to the formula *r* = log_2_(1 + 10^6^ · *x*) for each gene. Curated somatic mutations were retrieved from the Broad Institute TCGA GDAC Firehose Portal (http://gdac.broadinstitute.org/). Gene names were adapted to comply with those in the NCBI Entrez ID database as of July 7, 2015.

### Topological representations

We used the algorithm Mapper^28^, implemented in the Ayasdi software (https://www.ayasdi.com/platform/), to build topological representations of the RNA-seq data of each cancer cohort. Mapper builds upon any dimensional reduction algorithm (also known as “filter function”) to produce a new low-dimensional network representation on which local relationships are preserved. To that end, Mapper covers the low dimensional representation with overlapping bins and performs single-linkage clustering of the points in the high-dimensional space. The number of bins and their overlap are specified by the “resolution” and “gain” parameters respectively. The number of clusters in each bin is determined by the method described in Singh *et al*.^28^. A low-dimensional network is then built by assigning a node to each cluster, and if a sample appears in two nodes they are connected by an edge. A more detailed description of the Mapper algorithm for biologists can be found in the Methods section and Supplementary Note of Rizvi *et al*.^25^.

The output of Mapper is sensitive to several algorithmic choices. In our application, the following choices were made:

– *Metric.* We used Pearson’s correlation distance using the top 4,500 genes with highest variance as a measure of the similarity among the expression profile of tumors. We did not observed substantial differences between using Pearson’s and Spearman’s correlation distance in our analyses. We therefore used Pearson’s correlation distance given its reduced computation time in large datasets.
– *Filter function.* We built a *k* = 30 nearest neighbors graph using Pearson’s correlation distances between the samples and used a 2-dimensional embedding of the shortest path distances on this graph as the filter function. This choice filter function was based on the ability to capture biological proxies, such as the separation between the expression profiles of normal and tumor samples and the identification of known driver genes. Other choices of 2-dimensional filter functions, such as Principal Component Analysis or Multidimensional Scaling led to consistent results.
– *“Resolution” and “gain” parameters.* We covered the low-dimensional representation with overlapping squared bins. We scanned across the entire “resolution” and “gain” parameter space of the cover, as described in the paragraph “*Parameter scan and selection*”, obtaining stable results.

### Statistical analysis

We used the notions of topological association introduced in Rizvi *et al*.^25^ to identify features associated to localized regions of a phenotypic space. Our approach is closely related to the Laplacian score of He, Cai, and Niyogi^55,56^, and complementary to other statistical methods for network analysis^57^. In our case, the features that we tested were the somatic mutations in the tumor cohort, and the phenotypic space was the expression space of the tumor cohort. More specifically, for each mutated gene *g* in the cohort, we defined the following score:

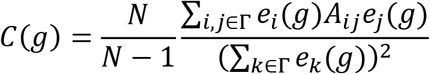

where Γ denotes the set of nodes in the topological representation, *A_ij_* its adjacency matrix, *N* the number of nodes in the representation, and *e_i_*(*g*) the average frequency of non-synonymous mutations of *g* for the samples in node *i*. The score *C*(*g*) is therefore a sum over the edges of the network, where the contribution of each edge is proportional to the product of the fraction of tumors that harbor the mutated gene in each of the two nodes connected by the edge. To be able to compare the score of mutated genes with different prevalence, we introduced a permutation test for each gene. A null distribution was built for *C*(*g*) by randomly permuting the patient id’s in the exome data and a *p*-value was assigned to the score of each gene *g* according to its null distribution. We performed 10^4^ permutations to build the null distribution of each gene. We controlled the false discovery rate (FDR) using the Benjamini-Hochberg (BH) procedure^58^. To avoid too large corrections due to multiple hypothesis testing, we only considered mutated genes with a prevalence in the cohort above a given threshold. The thresholds used in each cohort are summarized in Supplementary Table 4. In addition, we limited the number of genes in each analysis to the 350 genes with highest ratio between non-synonymous and total number of mutations. These thresholds were empirically determined for each cohort by looking at the size of the BH correction that resulted at different choices of the thresholds. For some cancer types, avoiding a large BH correction required relatively stringent thresholds (Supplementary Table 4), possibly reflecting noisier expression networks, e.g. due to differences in tumor purity among patients.

### Parameter scan and selection

To optimize the sensitivity of our approach at a fixed false discovery rate and control for the stability of the results against parameter choices, we generated 49 topological representations for the expression data of each tumor type by scanning over the parameter space of the Mapper algorithm. The resolution parameter was taken in the range 10 to 80, in intervals of 10, and the gain parameter 1.5-8.5, in intervals of 1. For each topological network, the statistical analysis performed in the previous paragraph was performed independently. We then selected a finer region in the parameter space for each cohort based on the following criteria:

– A large number of mutated genes with a significant score (*q*-value < 0.15) at a fixed FDR.
– Absence of significant spurious correlations and batch effects (as described in next paragraph).

For each selected region in the parameter space, we performed a finer scan across the resolution and gain parameters, taking intervals of 5 and 0.5 respectively.

### Control of spurious associations with expression

Hypermutated tumors often have a distinctive expression signature. In those cases, some localized regions of the expression space will consist of tumors with a higher mutation rate.

Those localized regions will harbor an accumulation of passenger mutations that may confound our approach. To control for associations between global expression patterns and the tumor mutation rate, we assessed the localization of the mutational tumor burden on the topological representations using the same approach as described in the paragraph “*Statistical analysis”*, where *e_i_* is now the average frequency of somatic mutations for the samples in node *i*. If the localization of the mutational burden was significant (*p*-value < 0.05), we manually set a threshold on the mutational burden to split the cohort into hypermutated and non-hypermutated tumors. This process could have been automated, however we found it unnecessary as small changes in the threshold do not affect substantially the results. The thresholds used in each cohort are summarized in Supplementary Table 4. We randomly subsampled mutations from each of the hypermutated tumors so that after subsampling the median mutational burden for hypermutated tumors in the cohort was equal to the median mutational burden for non-hypermutated tumors. We reassessed the significance of the localization of the mutational burden using the down sampled data. If the degree of localization of the mutational tumor burden was not significant, we continued the analysis using the down sampled mutation data. Otherwise, if the degree of localization was still significant after subsampling, we did not include the cohort in our study.

To control for associations between expression and mutation rates within the same gene, such as those due to transcription-coupled DNA repair, we assessed the similarity between the profiles of somatic mutation and mRNA expression on the topological representations. To that end, we computed the Jensen-Shannon divergence between the expression and mutation profiles of each gene in the topological representations using the formula

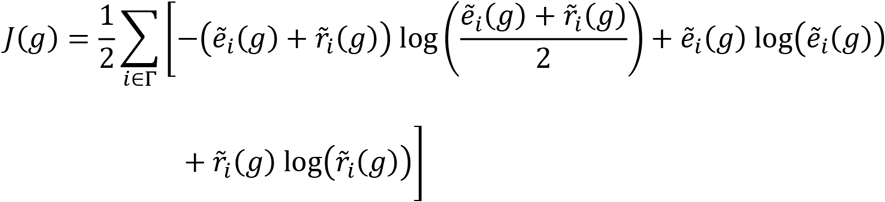

where 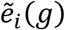 and 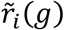 are respectively the fraction of tumors with gene *g* somatically mutated and the average expression of gene *g* in the tumors associated to the *i*-th node of the topological representation, normalized such that

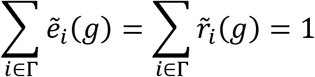

The significance of *J*(*g*) was assessed for each gene independently by means of a permutation test. To that end, for each gene the labels of the samples on the mutation data were randomly permuted 2,000 times, and *J*(*g*) was computed in each permutation. A *p*-value was estimated by counting the fraction of permutations that led to a value of *J*(*g*) smaller than the original value. Genes with a *p*-value for *J*(*g*) closed to 0 displayed a large degree of correlation between expression and mutation in the topological representation, whereas genes with a *p*-value for *J*(*g*) closed to 1 displayed a large degree of anticorrelation between expression and mutation in the topological representation. After adjusting for multiple hypothesis testing using Benjamini-Hochberg procedure to control the false discovery rate, we removed genes from the analysis for which the median *q*-value for *J*(*g*) across the parameter space of the topological representation was above 0.8, as those are potentially related to spurious anti-correlations between gene expression and mutation.

Last, to control for the presence of batch effects due to differences among mutation calling centers, we assessed the degree of localization of batches in the topological representation using the same approach as described in the paragraph *“Statistical analysis”,* with *e_i_* now represents the fraction of tumors in node *i* that were processed by a given center. We removed the contribution of batches whose degree of localization was significant (*p*-value < 0.05) according to this procedure.

### Computational benchmarking

We generated smaller LGG datasets by randomly sampling 50, 100, 200, 300, and 400 patients from the original LGG cohort. For each of these sizes, we generated a null data set by randomly permuting the labels of the patients on the expression data. We ran the integrative topological analysis in each of these new data sets using the same parameters than in the original analysis of the LGG cohort (Supplementary Table 4).

To benchmark the performance of algorithms based on a gold-standard list of cancer-associated genes, we followed the same approach as in Bertrand *et al*.^35^. We considered the same gold-standard list as in that reference. For each of the algorithms evaluated, we computed the precision (P), recall (R), and F1 score based the top min(15, *G*) significant genes (*q*-value < 0. 15)

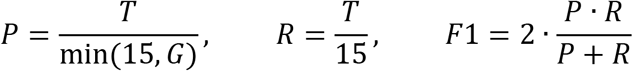

where *G* is the total number of significant genes and *T* the number of top min(15, *G*) significant genes present in the gold-standard. We run Xseq, OncodriveFML and 20/20+ with default parameters, as described in their documentation.

### MutSig2CV analyses

We downloaded from the Broad Institute TCGA GDAC Firehose Portal (http://gdac.broadinstitute.org/) the MutSig2CV v3.1 analyses of each of the 12 TCGA cohorts (Supplementary Table 4).

### Online database

Representative topological representations and pre-computed statistics were deposited in an online database for each of the 12 tumor types considered in this study(https://rabadan.c2b2.columbia.edu/pancancer). The interface of the database allows to explore the results of the analysis interactively.

### Induction of lung tumors in mice

Mouse experiments were performed following the institutional guidelines of the University of Oviedo (Comité de Ética en Experimentación Animal). *Adamts12^−/−^* mice were generated in a C57BL/6J genetic background and genotyped as in El Hour *et al*.^49^. Lung tumors were induced in 6-8 weeks old mice by intraperitoneal injection of 8 doses of 1 g/kg of urethane (ethyl carbamate; Sigma); second dose was given 48h after the initial one and then once a week to reach a total of 8 doses. Mice were sacrificed 20 weeks after the first urethane injection and during this time mice were fed *ad libitum*. Left lungs were fixed in 4% paraformaldehyde, paraffin-embedded and sectioned every 100 μm in of 10 μm slices. These were then stained with hematoxylin/eosin for morphological examination by experienced pathologists (Unidad de Histopatología Molecular en Modelos Animales de Cáncer, IUOPA). Tumors were quantified and classified according to their diameter in large (> 400 μm), medium (200-400 μm) and small (<200 μm) tumors.

### Generation of shADAMTS12 LL/2-luc-M38 cells

We used an *Adamts12* Mouse shRNA Plasmid (OriGene, Locus ID: 239337) and transfected LLC/2-luc-M38 (Caliper) cells with lipofectamine/plus (ThermoFisher Scientific) following the recommendations of manufacturer. We checked transfected cells for ADAMTS12 expression by western-blot for ADAMTS12 (Santa Cruz Biotechnology H-142) and β-actin (Sigma-Aldrich AC-15) in 10% polyacrylamide gels. Immunoreactive proteins were visualized using HRP-peroxidase-labeled anti-rabbit or anti-mouse secondary antibodies and the ECL detection system (Pierce).

### Proliferation assay

Cell proliferation was measured using the CellTiter 96 Non-radiactive Cell Proliferation Assay kit (Promega). LL/2-luc-M38 cells (3×10^4^/well) were seeded into 96-well plates in six replicates. Cell proliferation rates were determined on five consecutive days using the automated microtiter plate reader Power Wave WS (BioTek).

### Invasion assay

In-vitro invasion potential was assessed using 24-well Matrigel-coated invasion chambers with 8 μm pore size (BD Biosciences). A total of 5×10^4^ cells were allowed to migrate for 24 h using 10 % fetal bovine serum as chemoattractant. Cells that reached the lower surface were stained with crystal violet. At least three independent experiments were performed with triplicates for each condition. Cells were counted in 8 randomly selected microscopic fields.

### Immunohistochemistry

Lungs were fixed in 4% formalin for 24 h. After fixation, samples were dehydrated and embedded in paraffin. Sections of 4-μm thick were stained with hematoxylin and eosin for microscopy examination and consecutive sections were used for immunohistochemical labeling. Sections were incubated with anti-ADAMTS12 (H-142, Santa Cruz Biotechnologies, 1h at 37 °C) or with anti-Ki67 (ab66155, Abcam, o/n at 4°C) primary antibodies. Sections were then incubated 30 minutes with EnVision™+/HRP (Dako) and 5 minutes with Liquid DAB (Dako). Samples were counterstained with hematoxilin.

### Code availability

The source code and scripts used in the paper have been deposited in GitHub (https://github.com/CamaraLab/TDA-TCGA/).

### Author contributions

P.G.C. and R.R. conceived the methodology. U.R. implemented the computational pipeline and built the topological representations. Y.M., S.C., and A.J.O. conceived and performed the experiments. T.C. and O.E. designed and implemented the online database. A.J.L. and L.A. contributed to the analysis. All authors discussed the results and implications and collaborated on the writing of the manuscript.

## Acknowledgments

We thank J. A. Vega and IUOPA (Servicio de Histopatología Molecular en Modelos Animales de Cáncer) for assistance with the histopathology of mouse tumors; R. Aubin, G. Carlsson, K. Govek, A. Iavarone, A. Lasorella, S. Lowe, F. Sanchez-Rivera, E. Troisi, and S. Woodhouse for scientific discussions; and S. Christen, D. Ramanan, I. Sagalovskiy, and S. Thuault-Restituito for technical and administrative support. R.R. acknowledges funding from NIH (U54 CA193313-01). PGC acknowledges funding from Stand-Up-To-Cancer Convergence 2.0.

