## Supplemental Figures and Tables for "Identification of Relevant Genetic Alterations in Cancer using Topological Data Analysis"

[†] Currently at Early Signal, New York.

### Supplementary Figures

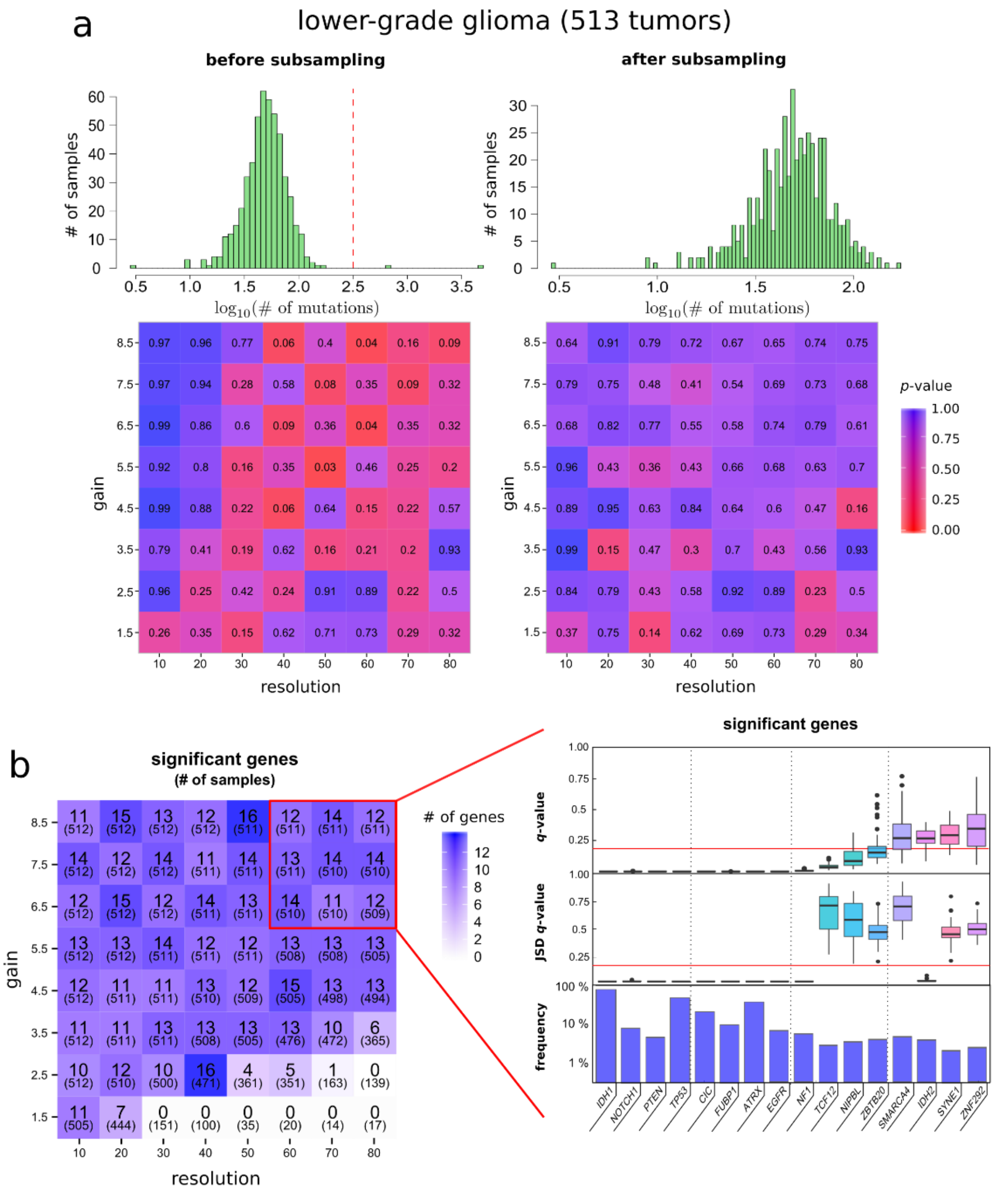


**Supplementary Figure 1 | Identification of mutated cancer genes in LGG using an integrative topological approach. (a)** Analysis of the localization of the mutational tumor burden in the reconstructed expression space of LGG. Top-left: histogram of the mutational burden for the 513 tumors in the LGG cohort. Note the logarithmic scale in the horizontal axis. The red dashed line separates hyper-mutated tumors ($n = 2$) from the rest. Bottom-left: statistical significance ($p$-value) of the localization of the mutational tumor burden in the topological representation, for different choices for the ‘gain’ and ‘resolution’ parameters of the Mapper algorithm. The mutational burden is significantly localized across a large region of the parameter space, indicating the presence of spurious correlations between the mutation rate and gene expression. Top-right: histogram of the mutational burden after down-sampling mutations in the hyper-mutated tumors. Bottom-right: statistical significance of the mutational tumor burden after down-sampling mutations in hyper-mutated tumors. The mutational burden is no-longer significantly localized across the parameter space of Mapper. (**b**) Left: number of significant genes at a fix false discovery rate ($q$-value < 0.15, Benjamini-Hochberg procedure), for different choices for the ‘gain’ and ‘resolution’ parameters of the Mapper algorithm. The number of samples in the largest connected component of the topological representation is indicated between parentheses. Right: list of mutated genes significantly localized in the reconstructed expression space of LGG for the region of the parameter space highlighted in red. From bottom to top, the prevalence of mutations in the cohort, the distribution of the statistical significance of the Jensen-Shannon distance between the expression and mutation profiles, and the distribution of the statistical significance of the localization in the topological representation across the region of the parameter space indicated in red, are displayed for each gene.

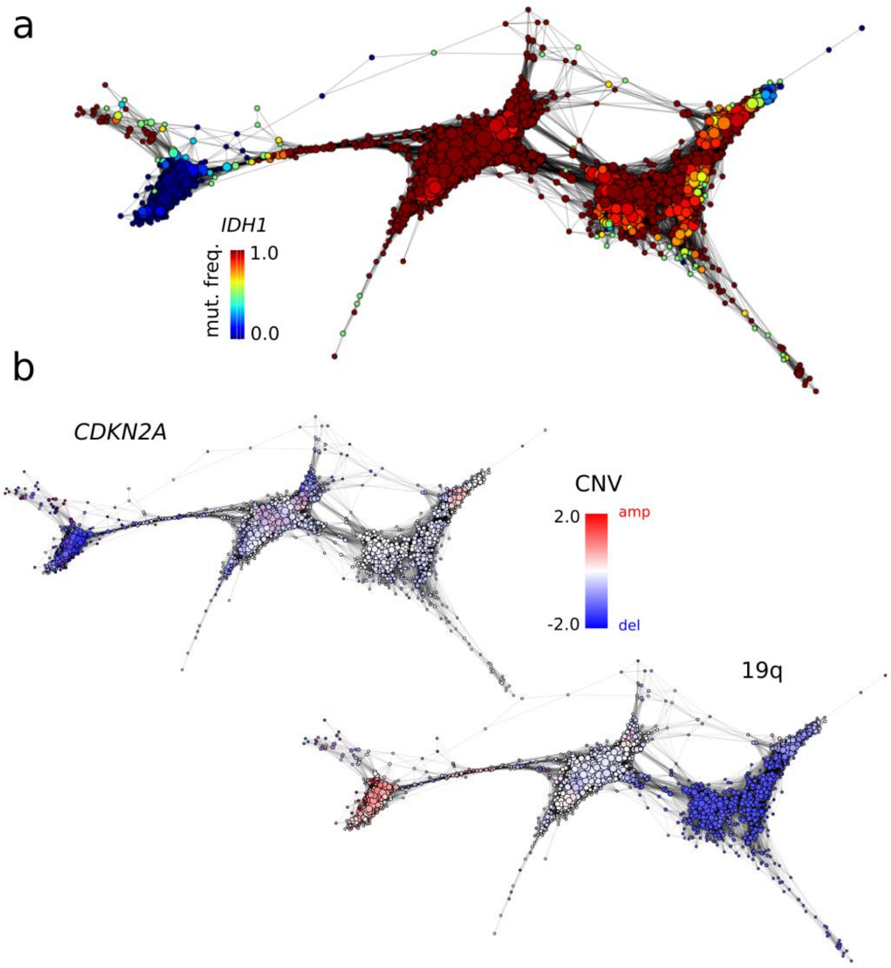


**Supplementary Figure 2 | Topological representation of the expression space LGG labeled according to the prevalence of various genomic alterations.** (**a**) Topological representation labeled by the prevalence of *IDH1* mutations. (**b**) Topological representation labeled by the prevalence of copy number alterations of the *CDKN2A* gene (top) and the chromosomal arm 19q (bottom).

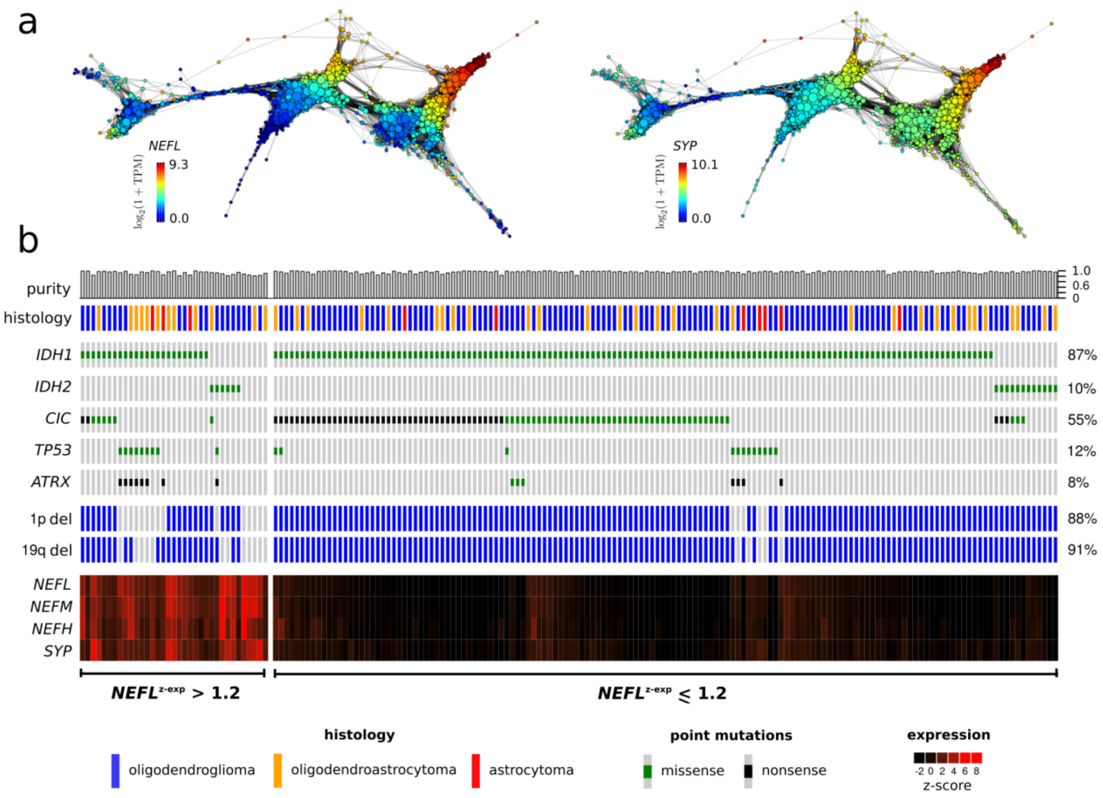


**Supplementary Figure 3 | Neuronal marker expression in LGG.** (**a**) Topological representation labeled by expression levels of *NEFL* (left) and *SYP* (right). (**b**) Co-mutation plot of the tumors in the oligodendrocytoma expression group. The purity estimates and histology type of the tumors is indicated. A heat map with the expression levels of various neuronal markers is also displayed. Tumors with high expression levels of neuronal markers harbor frequent deletions of the chromosome arm 19q, in addition to molecular alterations characteristic of astrocytic gliomas, such as *TP53* and *ATRX* mutations.

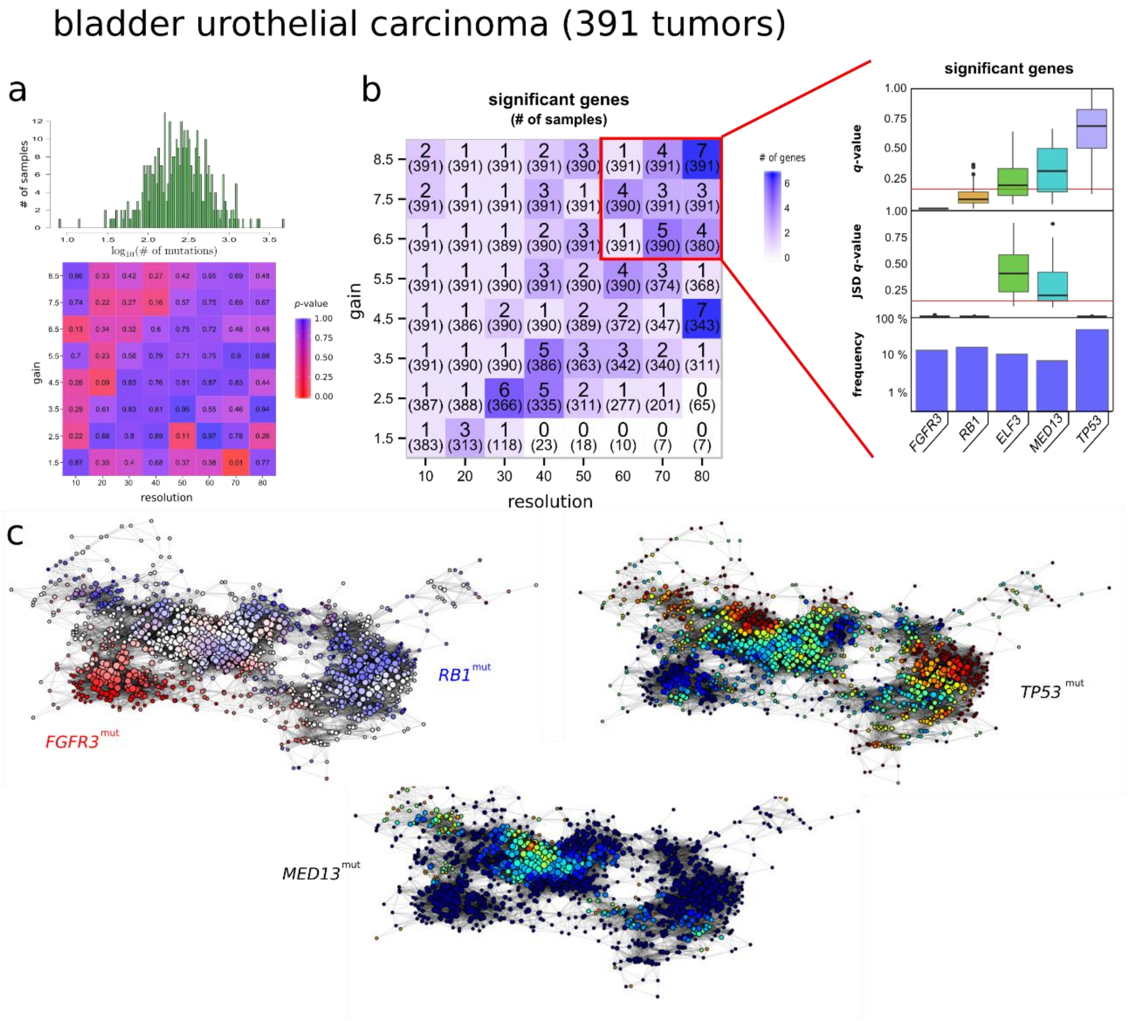


**Supplementary Figure 4 | Identification of mutated cancer genes in BLCA using an integrative topological approach.** (**a**) Analysis of the localization of the mutational tumor burden in the reconstructed expression space of BLCA. Top: histogram of the mutational burden for the 391 tumors in the BLCA cohort. Note the logarithmic scale in the horizontal axis. Bottom: statistical significance (*p*-value) of the localization of the mutational tumor burden in the topological representation, for different choices for the ‘gain’ and ‘resolution’ parameters of the

Mapper algorithm. The mutational burden is not significantly localized across the parameter space of Mapper. (**b**) Left: number of significant genes at a fix false discovery rate ($q$-value < 0.15, Benjamini-Hochberg procedure), for different choices for the 'gain' and 'resolution' parameters of the Mapper algorithm. The number of samples in the largest connected component of the topological representation is indicated between parentheses. Right: list of mutated genes significantly localized in the reconstructed expression space of BLCA for the region of the parameter space highlighted in red. From bottom to top, the prevalence of mutations in the cohort, the distribution of the statistical significance of the Jensen-Shannon distance between the expression and mutation profiles, and the distribution of the statistical significance of the localization in the topological representation across the region of the parameter space indicated in red, are displayed for each gene. (**c**) Topological representation labeled by the prevalence of mutations in the genes *FGFR3*, *RB1*, *TP53*, and *MED13*.

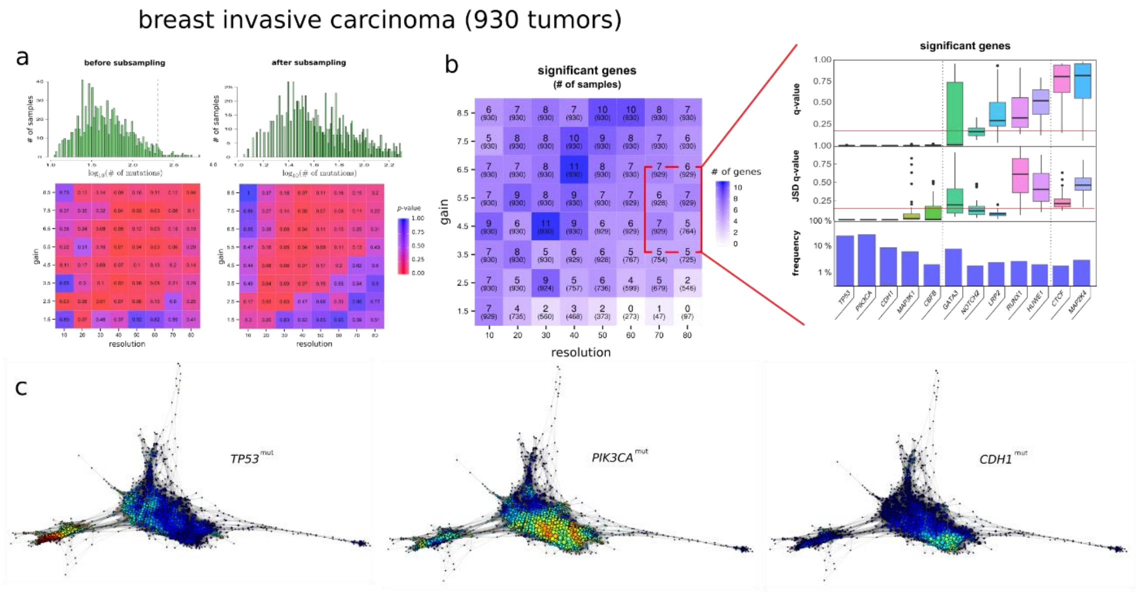


**Supplementary Figure 5 | Identification of mutated cancer genes in BRCA using an integrative topological approach.** (**a**) Analysis of the localization of the mutational tumor burden in the reconstructed expression space of BRCA. Top-left: histogram of the mutational burden for the 930 tumors in the BRCA cohort. Note the logarithmic scale in the horizontal axis. The red dashed line separates hyper-mutated tumors from the rest. Bottom-left: statistical significance (*p*-value) of the localization of the mutational tumor burden in the topological representation, for different choices for the ‘gain’ and ‘resolution’ parameters of the Mapper algorithm. The mutational burden is significantly localized across a large region of the parameter space, indicating the presence of spurious correlations between the mutation rate and gene expression. Top-right: histogram of the mutational burden after down-sampling mutations in the hyper-mutated tumors. Bottom-right: statistical significance of the mutational tumor burden after down-sampling mutations in hyper-mutated tumors. The mutational burden is no-longer significantly localized across the parameter space of Mapper. (**b**) Left: number of significant genes at a fix false discovery rate ($q$-value < 0.15, Benjamini-Hochberg procedure), for different choices for the 'gain' and 'resolution' parameters of the Mapper algorithm. The number of samples in the largest connected component of the topological representation is indicated between parentheses. Right: list of mutated genes significantly localized in the reconstructed expression space of BRCA for the region of the parameter space highlighted in red. From bottom to top, the prevalence of mutations in the cohort, the distribution of the statistical significance of the Jensen-Shannon distance between the expression and mutation profiles, and the distribution of the statistical significance of the localization in the topological representation across the region of the parameter space indicated in red, are displayed for each gene. (**c**) Topological representation labeled by the prevalence of mutations in the genes *TP53*, *PIK3CA*, and *CDH1*.

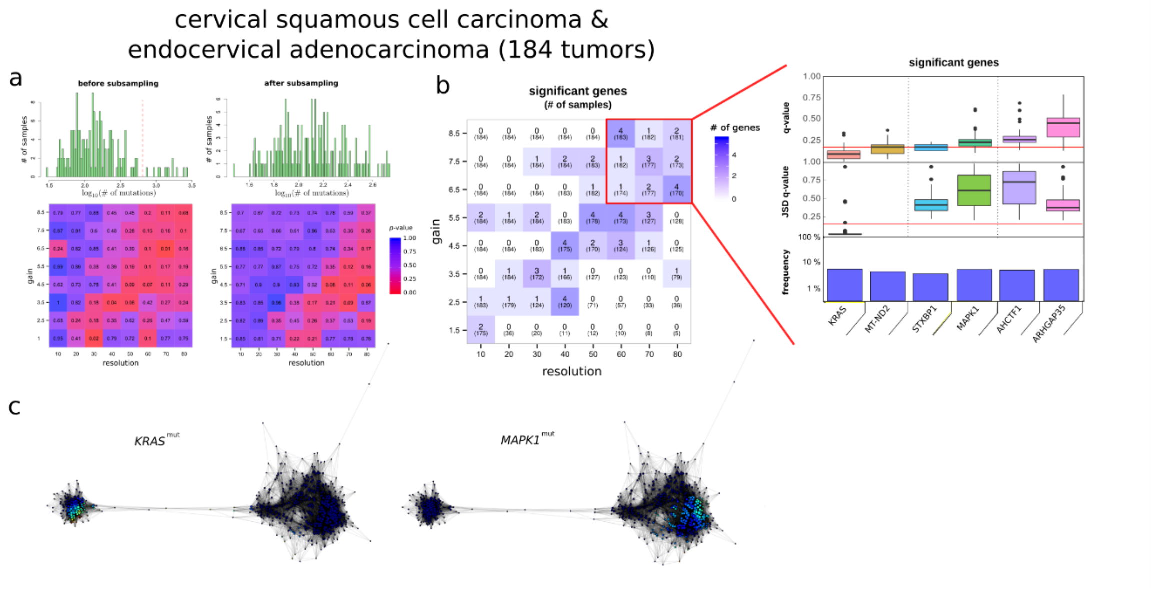


**Supplementary Figure 6 | Identification of mutated cancer genes in CESC using an integrative topological approach.** (**a**) Analysis of the localization of the mutational tumor burden in the reconstructed expression space of CESC. Top-left: histogram of the mutational burden for the 184 tumors in the CESC cohort. Note the logarithmic scale in the horizontal axis. The red dashed line separates hyper-mutated tumors from the rest. Bottom-left: statistical significance (*p*-value) of the localization of the mutational tumor burden in the topological representation, for different choices for the ‘gain’ and ‘resolution’ parameters of the Mapper algorithm. The mutational burden is significantly localized across a large region of the parameter space, indicating the presence of spurious correlations between the mutation rate and gene expression. Top-right: histogram of the mutational burden after down-sampling mutations in the hyper-mutated tumors. Bottom-right: statistical significance of the mutational tumor burden after down-sampling mutations in hyper-mutated tumors. The mutational burden is no-longer significantly localized across the parameter space of Mapper. (**b**) Left: number of significant genes at a fix false discovery rate ($q$-value < 0.15, Benjamini-Hochberg procedure), for different choices for the ‘gain’ and ‘resolution’ parameters of the Mapper algorithm. The number of samples in the largest connected component of the topological representation is indicated between parentheses. Right: list of mutated genes significantly localized in the reconstructed expression space of CESC for the region of the parameter space highlighted in red. From bottom to top, the prevalence of mutations in the cohort, the distribution of the statistical significance of the Jensen-Shannon distance between the expression and mutation profiles, and the distribution of the statistical significance of the localization in the topological representation across the region of the parameter space indicated in red, are displayed for each gene. (**c**) Topological representation labeled by the prevalence of mutations in the genes *KRAS* and *MAPK1*.

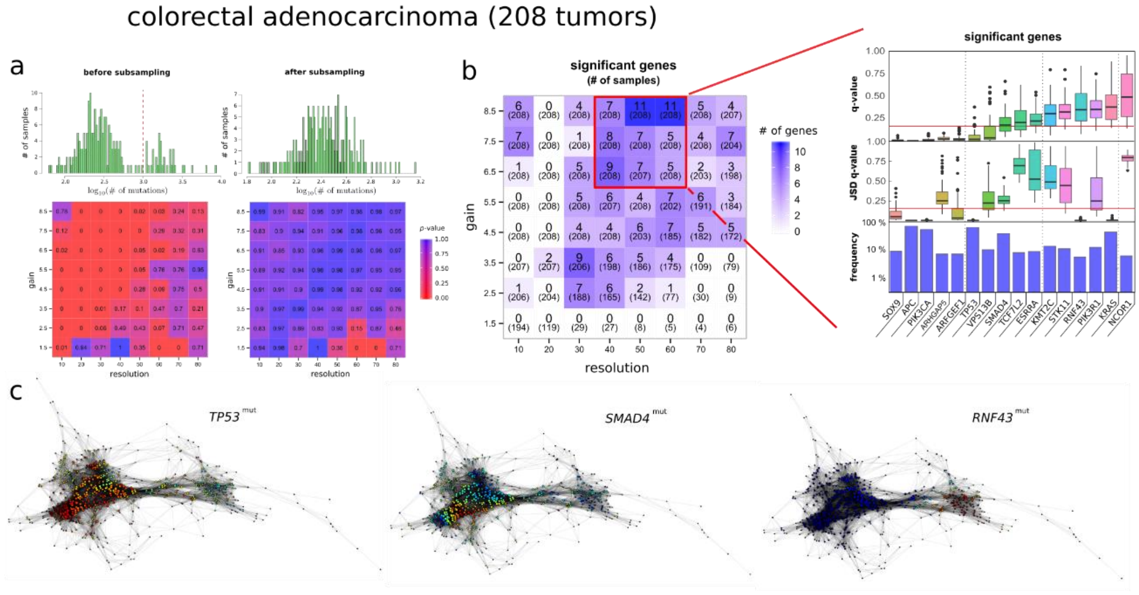


**Supplementary Figure 7 | Identification of mutated cancer genes in COAD using an integrative topological approach.** (**a**) Analysis of the localization of the mutational tumor burden in the reconstructed expression space of COAD. Top-left: histogram of the mutational burden for the 208 tumors in the COAD cohort. Note the logarithmic scale in the horizontal axis. The red dashed line separates hyper-mutated tumors from the rest. Bottom-left: statistical significance (*p*-value) of the localization of the mutational tumor burden in the topological representation, for different choices for the ‘gain’ and ‘resolution’ parameters of the Mapper algorithm. The mutational burden is significantly localized across a large region of the parameter space, indicating the presence of spurious correlations between the mutation rate and gene expression. Top-right: histogram of the mutational burden after down-sampling mutations in the hyper-mutated tumors. Bottom-right: statistical significance of the mutational tumor burden after down-sampling mutations in hyper-mutated tumors. The mutational burden is no-longer significantly localized across the parameter space of Mapper. (**b**) Left: number of significant genes at a fix false discovery rate ($q$-value < 0.15, Benjamini-Hochberg procedure), for different choices for the 'gain' and 'resolution' parameters of the Mapper algorithm. The number of samples in the largest connected component of the topological representation is indicated between parentheses. Right: list of mutated genes significantly localized in the reconstructed expression space of COAD for the region of the parameter space highlighted in red. From bottom to top, the prevalence of mutations in the cohort, the distribution of the statistical significance of the Jensen-Shannon distance between the expression and mutation profiles, and the distribution of the statistical significance of the localization in the topological representation across the region of the parameter space indicated in red, are displayed for each gene. (**c**) Topological representation labeled by the prevalence of mutations in the genes *TP53*, *SMAD4*, and *RNF43*.

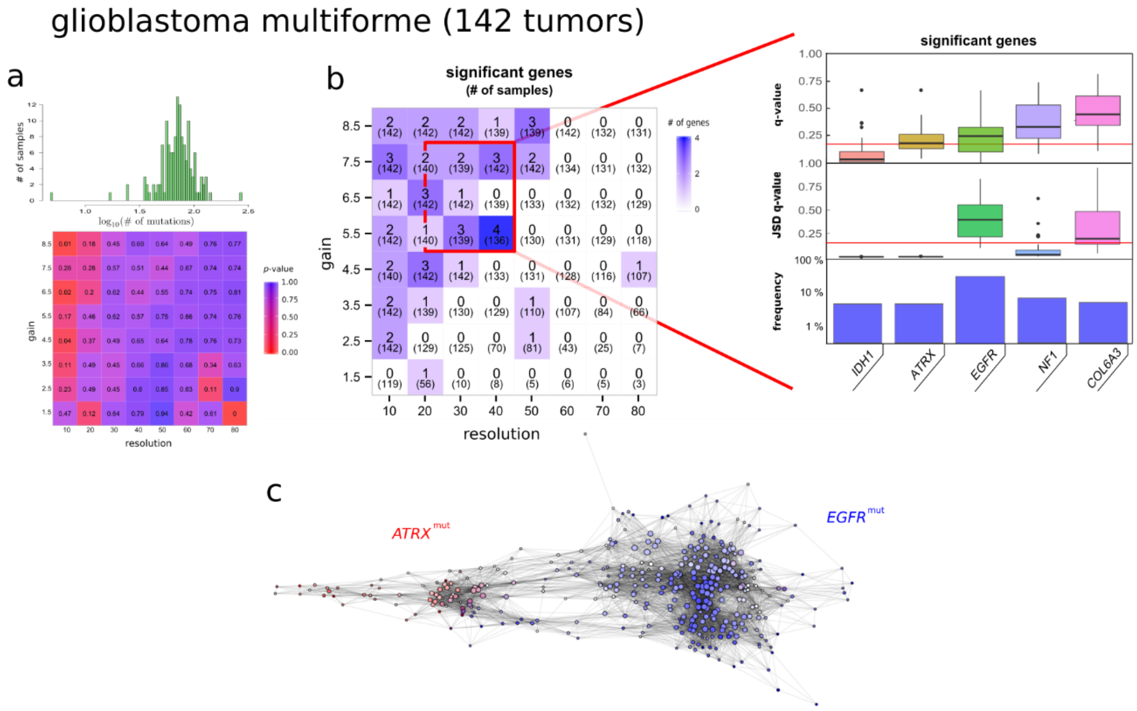


**Supplementary Figure 8 | Identification of mutated cancer genes in GBM using an integrative topological approach.** (**a**) Analysis of the localization of the mutational tumor burden in the reconstructed expression space of GBM. Top: histogram of the mutational burden for the 142 tumors in the GBM cohort. Note the logarithmic scale in the horizontal axis. Bottom: statistical significance (*p*-value) of the localization of the mutational tumor burden in the topological representation, for different choices for the 'gain' and 'resolution' parameters of the Mapper algorithm. The mutational burden is not significantly localized across the parameter space of Mapper. (**b**) Left: number of significant genes at a fix false discovery rate ($q$-value < 0.15, Benjamini-Hochberg procedure), for different choices for the 'gain' and 'resolution' parameters of the Mapper algorithm. The number of samples in the largest connected component of the topological representation is indicated between parentheses. Right: list of mutated genes significantly localized in the reconstructed expression space of GBM for the region of the parameter space highlighted in red. From bottom to top, the prevalence of mutations in the cohort, the distribution of the statistical significance of the Jensen-Shannon distance between the expression and mutation profiles, and the distribution of the statistical significance of the localization in the topological representation across the region of the parameter space indicated in red, are displayed for each gene. (**c**) Topological representation labeled by the prevalence of mutations in the genes *ATRX* and *EGFR*.

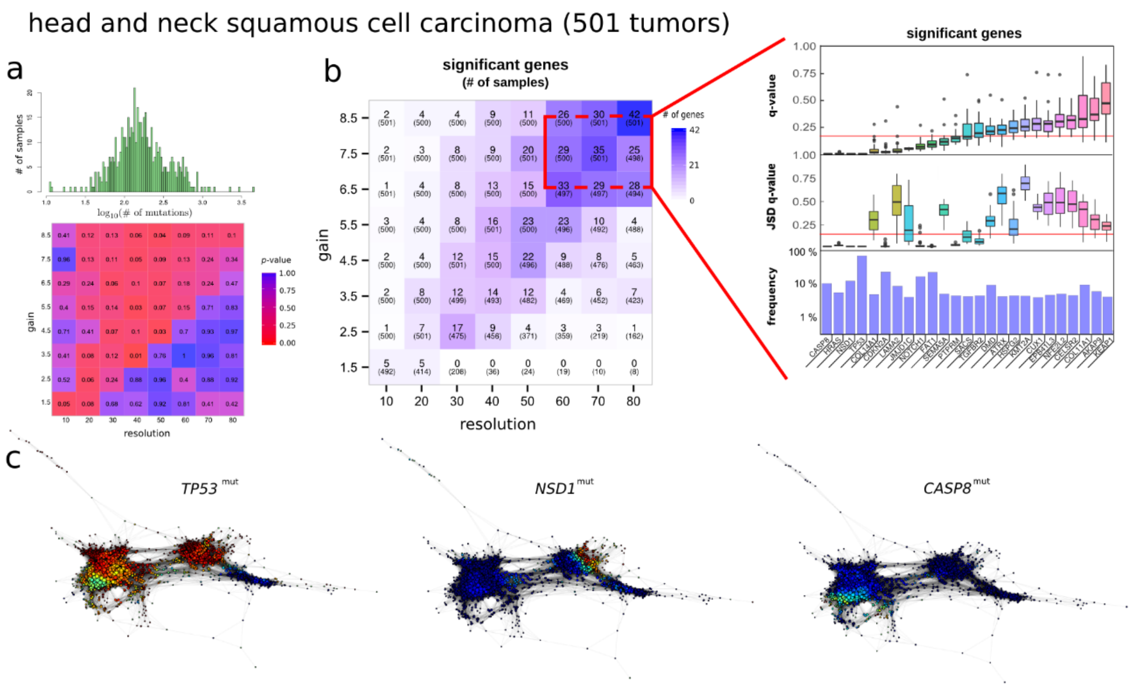


**Supplementary Figure 9 | Identification of mutated cancer genes in HNSC using an integrative topological approach.** (**a**) Analysis of the localization of the mutational tumor burden in the reconstructed expression space of HNSC. Top: histogram of the mutational burden for the 501 tumors in the HNSC cohort. Note the logarithmic scale in the horizontal axis. Bottom: statistical significance ($p$-value) of the localization of the mutational tumor burden in the topological representation, for different choices for the 'gain' and 'resolution' parameters of the Mapper algorithm. The mutational burden is not significantly localized across the parameter space of Mapper. (**b**) Left: number of significant genes at a fix false discovery rate ($q$-value < 0.15, Benjamini-Hochberg procedure), for different choices for the 'gain' and 'resolution' parameters of the Mapper algorithm. The number of samples in the largest connected component of the topological representation is indicated between parentheses. Right: list of mutated genes significantly localized in the reconstructed expression space of HNSC for the region of the parameter space highlighted in red. From bottom to top, the prevalence of mutations in the cohort, the distribution of the statistical significance of the Jensen-Shannon distance between the expression and mutation profiles, and the distribution of the statistical significance of the localization in the topological representation across the region of the parameter space indicated in red, are displayed for each gene. (**c**) Topological representation labeled by the prevalence of mutations in the genes *TP53*, *NSD1*, and *CASP8*.

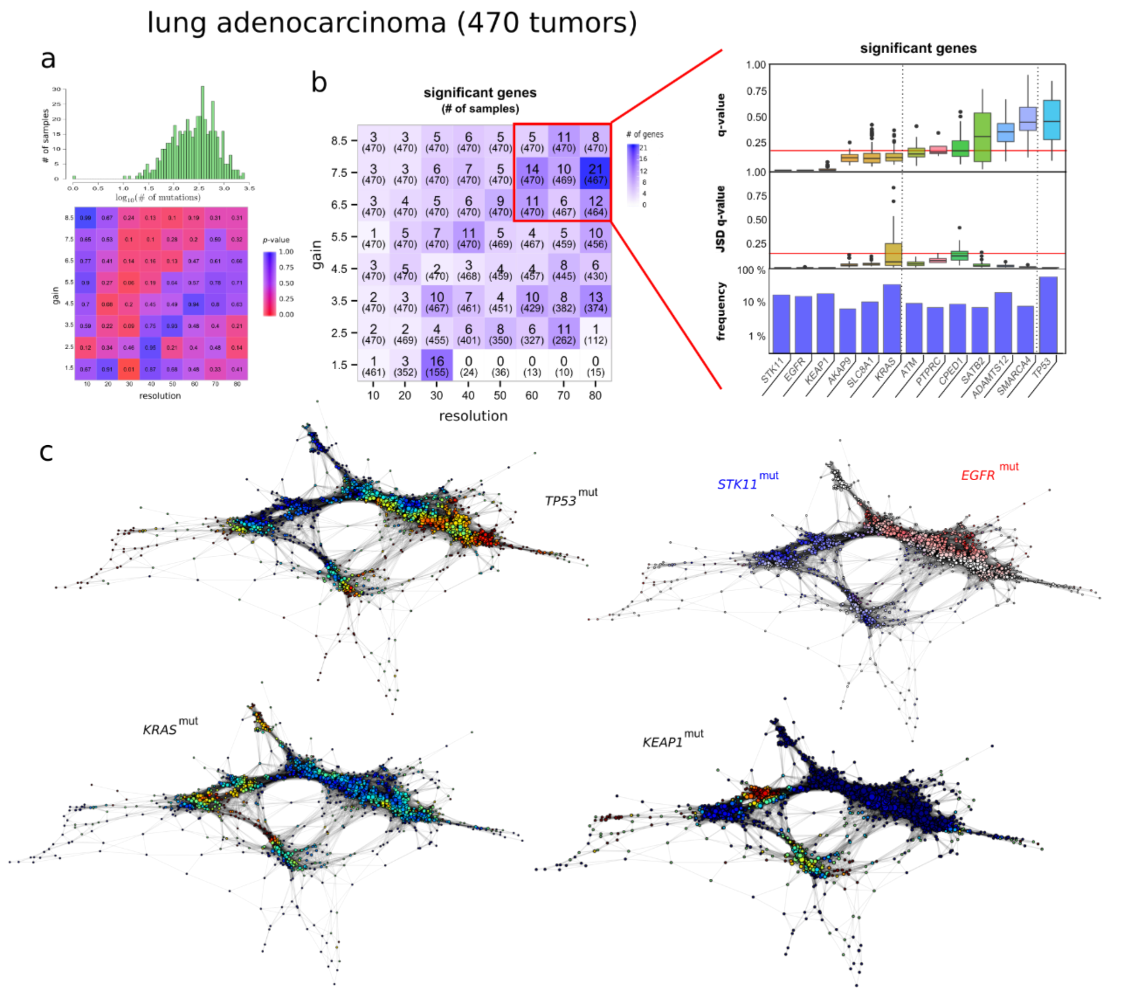


**Supplementary Figure 10 | Identification of mutated cancer genes in LUAD using an integrative topological approach.** (**a**) Analysis of the localization of the mutational tumor burden in the reconstructed expression space of LUAD. Top: histogram of the mutational burden for the 470 tumors in the LUAD cohort. Note the logarithmic scale in the horizontal axis. Bottom: statistical significance (*p*-value) of the localization of the mutational tumor burden in the topological representation, for different choices for the ‘gain’ and ‘resolution’ parameters of the Mapper algorithm. The mutational burden is not significantly localized across the parameter space of Mapper. (**b**) Left: number of significant genes at a fix false discovery rate (*q*-value < 0.15, Benjamini-Hochberg procedure), for different choices for the 'gain' and 'resolution' parameters of the Mapper algorithm. The number of samples in the largest connected component of the topological representation is indicated between parentheses. Right: list of mutated genes significantly localized in the reconstructed expression space of LUAD for the region of the parameter space highlighted in red. From bottom to top, the prevalence of mutations in the cohort, the distribution of the statistical significance of the Jensen-Shannon distance between the expression and mutation profiles, and the distribution of the statistical significance of the localization in the topological representation across the region of the parameter space indicated in red, are displayed for each gene. (**c**) Topological representation labeled by the prevalence of mutations in the genes *TP53, STK11, EGFR, KRAS*, and *KEAP1*.

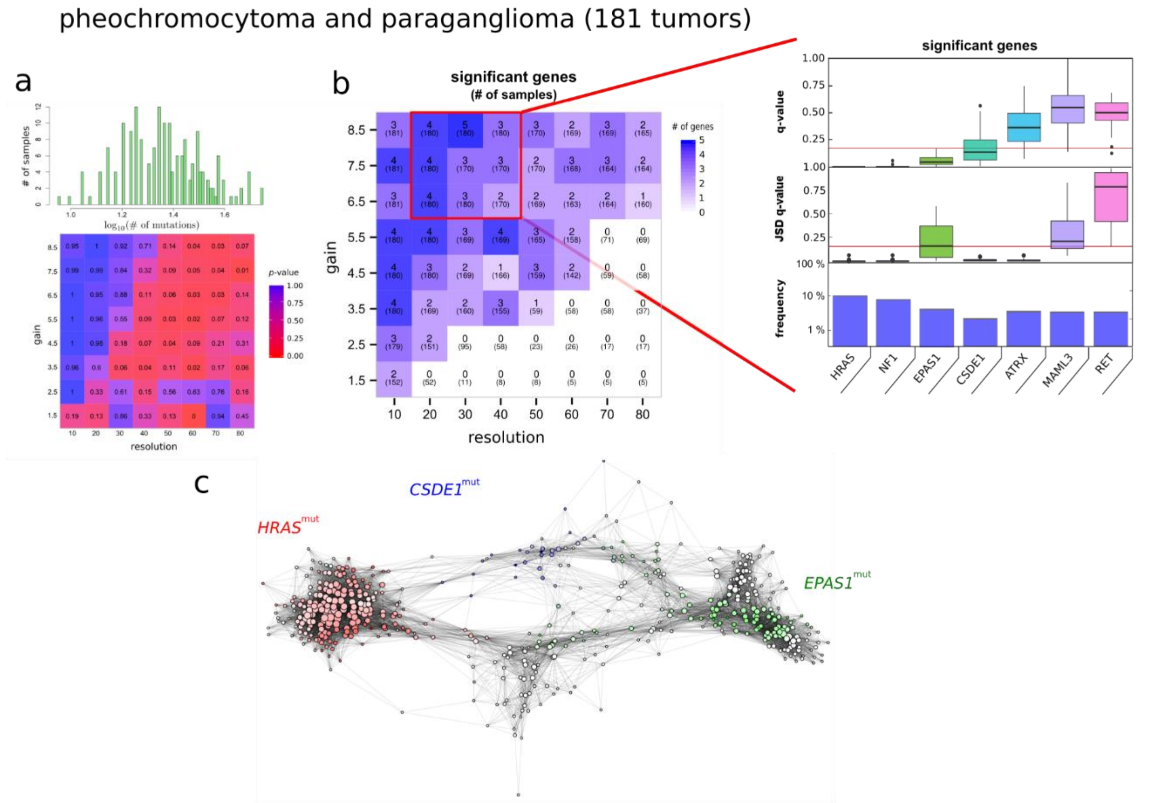


**Supplementary Figure 11 | Identification of mutated cancer genes in PCPG using an integrative topological approach.** (**a**) Analysis of the localization of the mutational tumor burden in the reconstructed expression space of PCPG. Top: histogram of the mutational burden for the 181 tumors in the PCPG cohort. Note the logarithmic scale in the horizontal axis. Bottom: statistical significance (*p*-value) of the localization of the mutational tumor burden in the topological representation, for different choices for the ‘gain’ and ‘resolution’ parameters of the Mapper algorithm. The mutational burden is not significantly localized across the parameter space of Mapper. (**b**) Left: number of significant genes at a fix false discovery rate ($q$-value < 0.15, Benjamini-Hochberg procedure), for different choices for the ‘gain’ and ‘resolution’ parameters of the Mapper algorithm. The number of samples in the largest connected component of the topological representation is indicated between parentheses. Right: list of mutated genes significantly localized in the reconstructed expression space of PCPG for the region of the parameter space highlighted in red. From bottom to top, the prevalence of mutations in the cohort, the distribution of the statistical significance of the Jensen-Shannon distance between the expression and mutation profiles, and the distribution of the statistical significance of the localization in the topological representation across the region of the parameter space indicated in red, are displayed for each gene. (**c**) Topological representation labeled by the prevalence of mutations in the genes *HRAS*, *EPAS1*, and *CSDE1*.

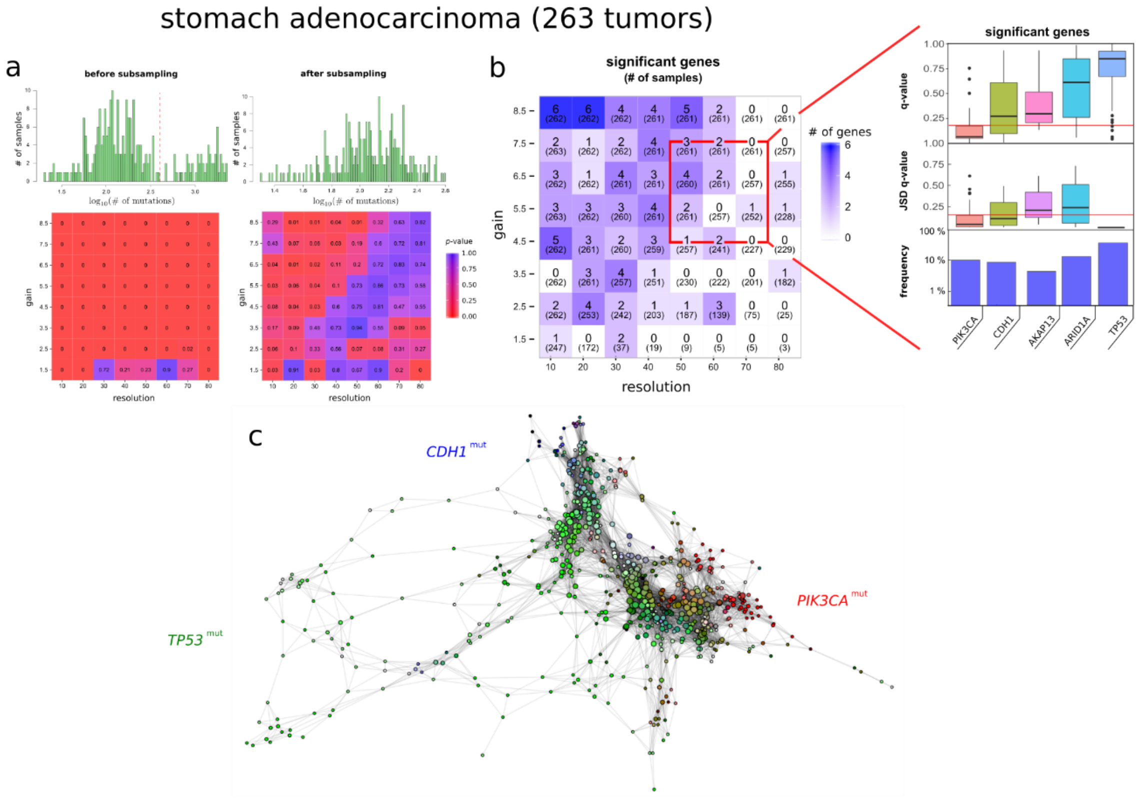


**Supplementary Figure 12 | Identification of mutated cancer genes in STAD using an integrative topological approach.** (**a**) Analysis of the localization of the mutational tumor burden in the reconstructed expression space of STAD. Top: histogram of the mutational burden for the 263 tumors in the STAD cohort. Note the logarithmic scale in the horizontal axis. Bottom: statistical significance (*p*-value) of the localization of the mutational tumor burden in the topological representation, for different choices for the 'gain' and 'resolution' parameters of the Mapper algorithm. The mutational burden is not significantly localized across the parameter space of Mapper. (**b**) Left: number of significant genes at a fix false discovery rate ($q$-value < 0.15, Benjamini-Hochberg procedure), for different choices for the 'gain' and 'resolution'

parameters of the Mapper algorithm. The number of samples in the largest connected component of the topological representation is indicated between parentheses. Right: list of mutated genes significantly localized in the reconstructed expression space of STAD for the region of the parameter space highlighted in red. From bottom to top, the prevalence of mutations in the cohort, the distribution of the statistical significance of the Jensen-Shannon distance between the expression and mutation profiles, and the distribution of the statistical significance of the localization in the topological representation across the region of the parameter space indicated in red, are displayed for each gene. (**c**) Topological representation labeled by the prevalence of mutations in the genes *TP53, CDH1,* and *PIK3CA.*

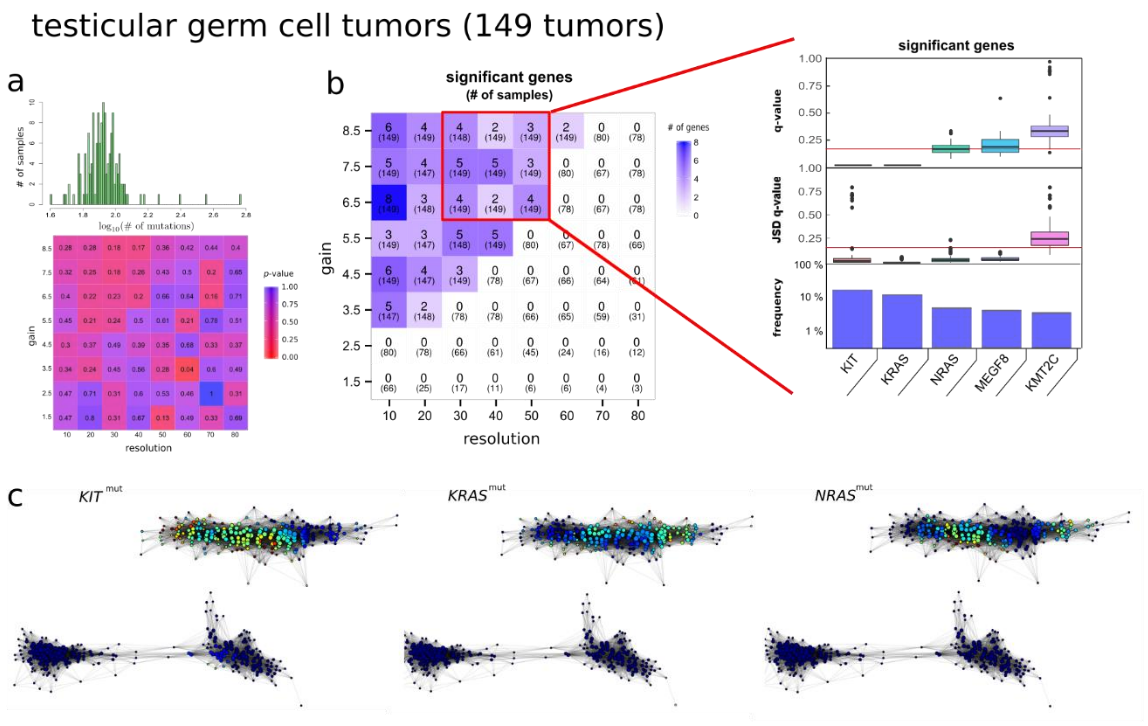


**Supplementary Figure 13 | Identification of mutated cancer genes in TGCT using an integrative topological approach.** (**a**) Analysis of the localization of the mutational tumor burden in the reconstructed expression space of TGCT. Top: histogram of the mutational burden for the 149 tumors in the TGCT cohort. Note the logarithmic scale in the horizontal axis. Bottom: statistical significance (*p*-value) of the localization of the mutational tumor burden in the topological representation, for different choices for the ‘gain’ and ‘resolution’ parameters of the Mapper algorithm. The mutational burden is not significantly localized across the parameter space of Mapper. (**b**) Left: number of significant genes at a fix false discovery rate ($q$-value < 0.15, Benjamini-Hochberg procedure), for different choices for the ‘gain’ and ‘resolution’ parameters of the Mapper algorithm. The number of samples in the largest connected component of the topological representation is indicated between parentheses. Right: list of mutated genes significantly localized in the reconstructed expression space of TGCT for the region of the parameter space highlighted in red. From bottom to top, the prevalence of mutations in the cohort, the distribution of the statistical significance of the Jensen-Shannon distance between the expression and mutation profiles, and the distribution of the statistical significance of the localization in the topological representation across the region of the parameter space indicated in red, are displayed for each gene. (**c**) Topological representation labeled by the prevalence of mutations in the genes *KIT*, *KRAS*, and *NRAS*.

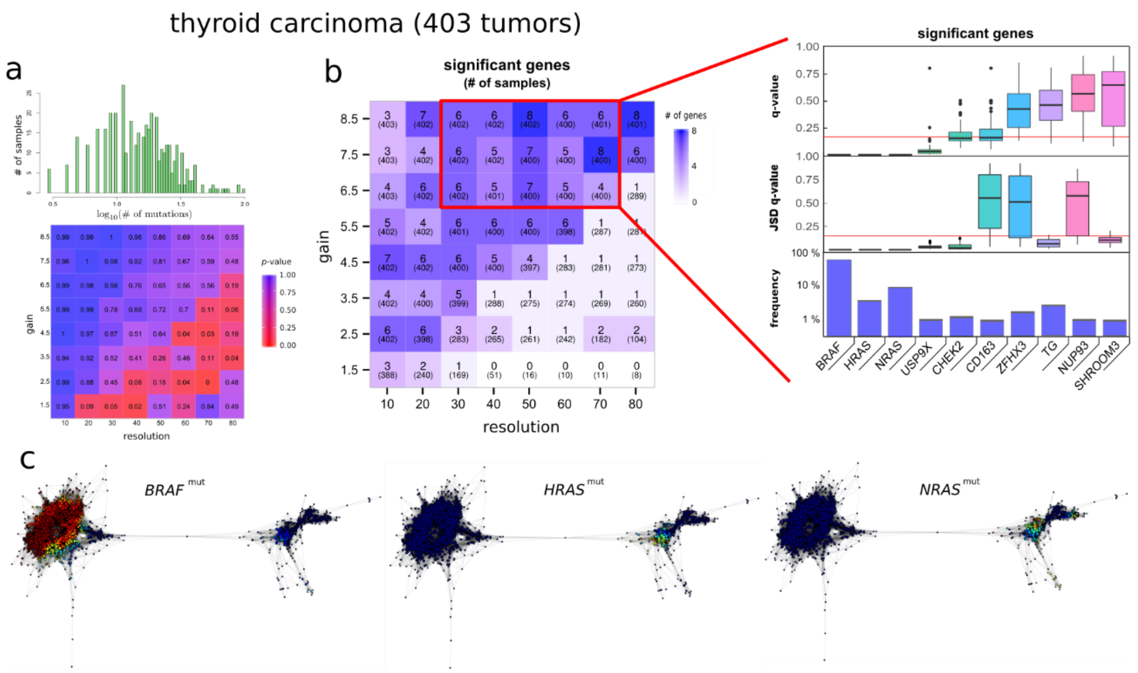


**Supplementary Figure 14 | Identification of mutated cancer genes in THCA using an integrative topological approach.** (**a**) Analysis of the localization of the mutational tumor burden in the reconstructed expression space of THCA. Top: histogram of the mutational burden for the 403 tumors in the THCA cohort. Note the logarithmic scale in the horizontal axis. Bottom: statistical significance (*p*-value) of the localization of the mutational tumor burden in the topological representation, for different choices for the 'gain' and 'resolution' parameters of the Mapper algorithm. The mutational burden is not significantly localized across the parameter space of Mapper. (**b**) Left: number of significant genes at a fix false discovery rate ($q$-value < 0.15, Benjamini-Hochberg procedure), for different choices for the 'gain' and 'resolution' parameters of the Mapper algorithm. The number of samples in the largest connected component of the topological representation is indicated between parentheses. Right: list of mutated genes significantly localized in the reconstructed expression space of THCA for the region of the parameter space highlighted in red. From bottom to top, the prevalence of mutations in the cohort, the distribution of the statistical significance of the Jensen-Shannon distance between the expression and mutation profiles, and the distribution of the statistical significance of the localization in the topological representation across the region of the parameter space indicated in red, are displayed for each gene. (**c**) Topological representation labeled by the prevalence of mutations in the genes *BRAF*, *HRAS*, and *NRAS*.

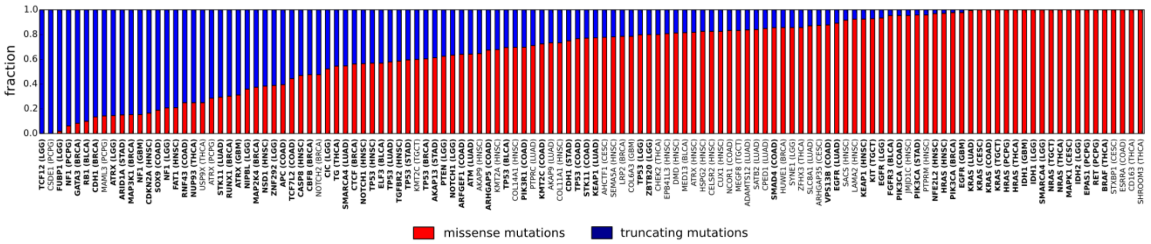


**Supplementary Figure 15 | Relative fraction of truncating and missense mutations for significant genes (*q*-value < 0.15) in the analysis of 12 tumor types.** Genes that were also significant (*q*-value < 0.15) under MutSig2CV are shown in bold.

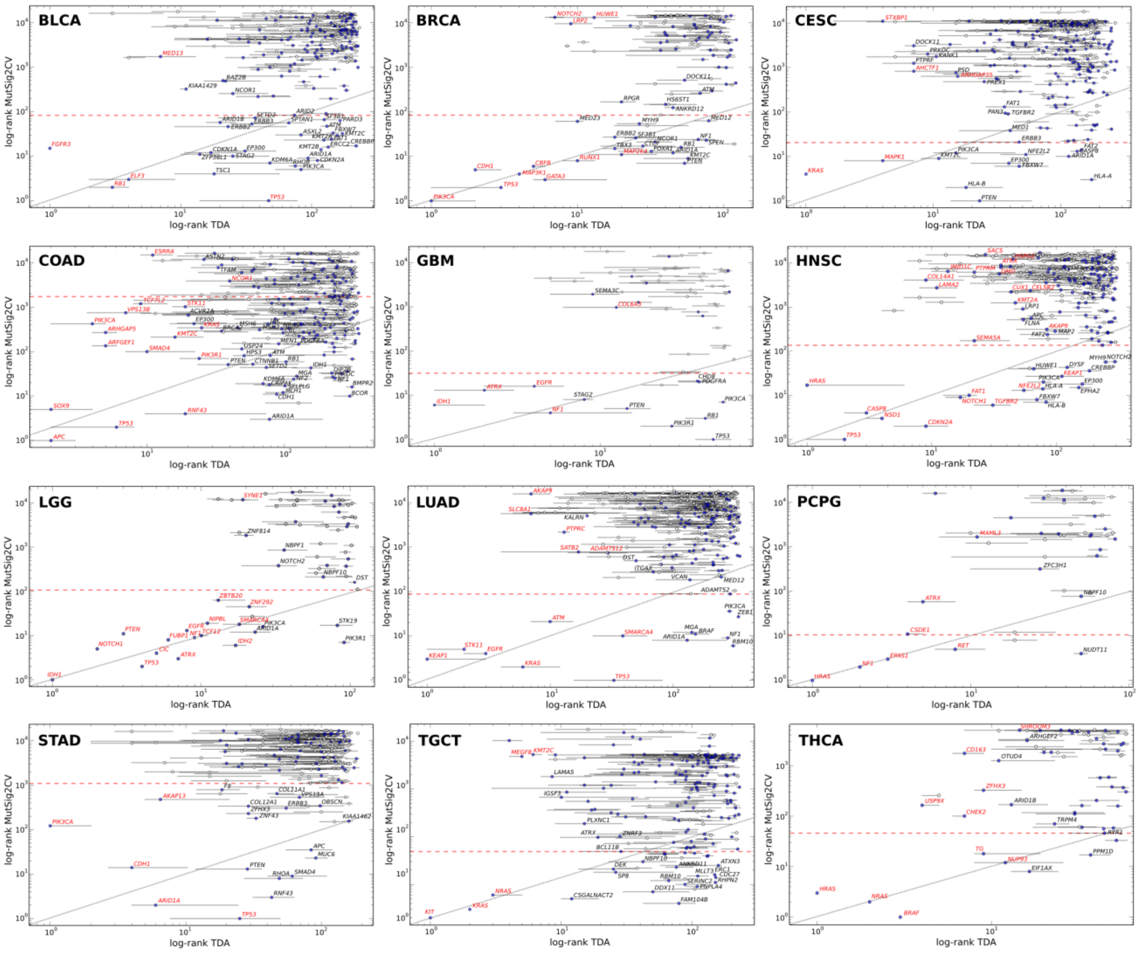


**Supplementary Figure 16 | Combination of the integrative topological and MutSig2CV analyses of 12 tumor types.** For each tumor type, the rank of significance according to the MutSig2CV analysis is plotted against the range of significance according to the integrative topological analysis, in log-log scale. Genes in red are significant (q-value < 0.15) in the integrative topological analysis. Genes below the red dashed line are significant (q-value < 0.15) in the MutSig2CV analysis. Horizontal bars indicate rank intervals containing 68% of the data across the selected parameter space of the Mapper algorithm (indicated by red squares in Supplementary Figs. 1 and 4-14). For reference, the diagonal is also shown in each plot.

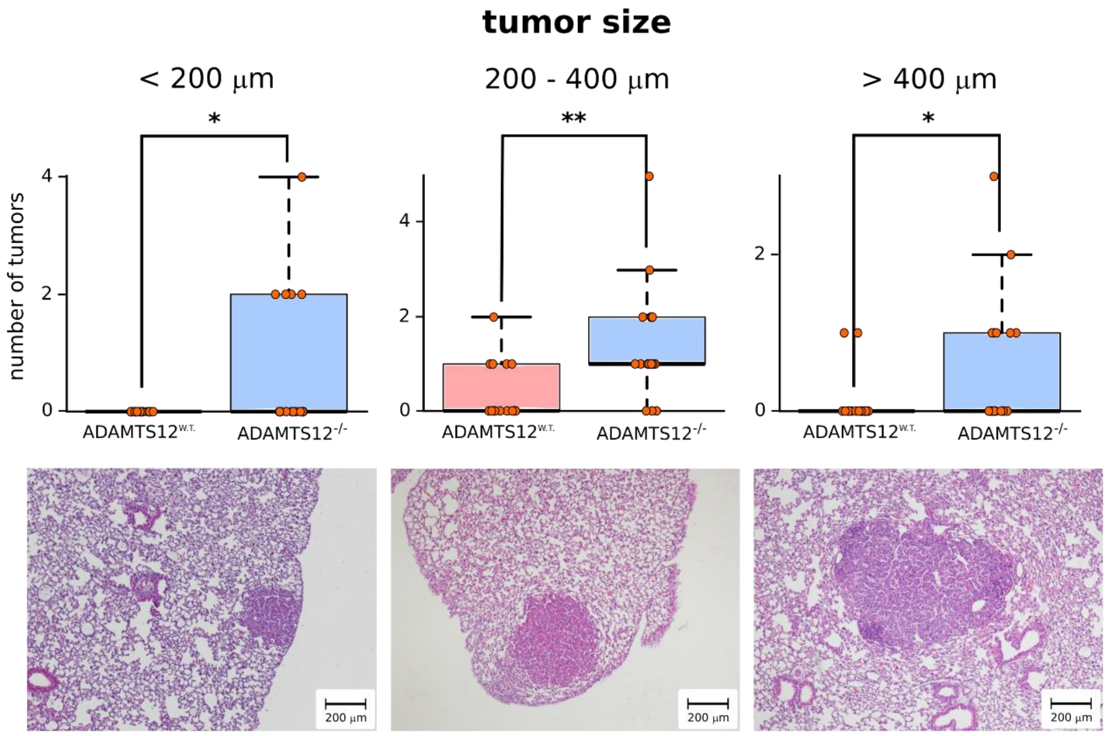


**Supplementary Figure 17 | Number of lung tumors induced by urethane in control and ADAMTS12-deficient mice disaggregated by tumor size.** Representative hematoxylin-eosin stained tissue sections displaying lung adenocarcinomas are displayed at the bottom. Statistical significances were determined using two-tailed Student's t-test ($p$ = 0.03 for tumors < 200 μm, $p$ = 0.004 for tumors between 200 and 400 μm, and $p$ = 0.05 for tumors > 400 μm).

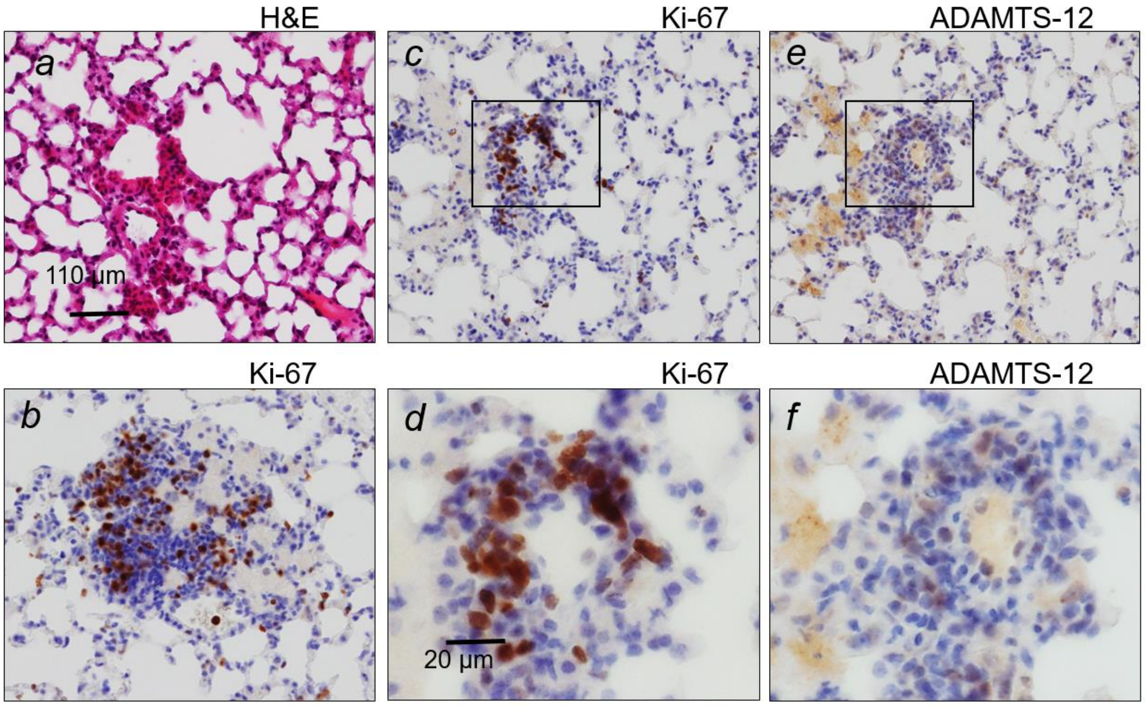


**Supplementary Figure 18 | Immunohistochemistry staining of urethane-induced lung tumors.** (**a**) Structural details and the identification of tumor areas were achieved by staining randomly selected sections with H&E. (**b**, **c** and **d**) Tumor areas showed strong nuclear staining for Ki-67. (**e** and **f**) Immunostaining for ADAMTS-12 was light but specific. In comparing approximate serial sections, no co-localization of Ki-67 and ADAMTS-12 was detected (c-d and e-f).

### Supplementary Tables

**Supplementary Table 1 | Comparison of the integrative topological approach to other algorithms for the identification of cancer-associated mutated genes.** For each algorithm, the precision (P), recall (R), and F1 score was computed based on the top 15 predictions in the LGG, COAD, and BRCA cohorts (see Methods). The best score in each columns is highlighted.

| | LGG | | | COAD | | | BRCA | | |
|---|---|---|---|---|---|---|---|---|---|
| **Algorithm** | **P** | **R** | **F1** | **P** | **R** | **F1** | **P** | **R** | **F1** |
| **TDA** | 0.87 | **0.87** | **0.87** | **0.67** | **0.67** | **0.67** | 0.92 | 0.73 | 0.81 |
| **MutSig2CV** | 0.87 | **0.87** | **0.87** | 0.53 | 0.53 | 0.53 | **0.93** | **0.93** | **0.93** |
| **Xseq** | **1.00** | 0.13 | 0.23 | - | - | - | - | - | - |
| **OncodriveFML** | 0.82 | 0.60 | 0.69 | 0.53 | 0.53 | 0.53 | 0.67 | 0.67 | 0.67 |
| **20/20+** | 0.61 | 0.53 | 0.57 | 0.40 | 0.40 | 0.40 | 0.53 | 0.53 | 0.53 |

**Supplementary Table 2 | Non-synonymous mutations for the significant genes in the integrative topological analysis of 12 tumor types.**

**[Provided as a separate file.]**

**Supplementary Table 3 | Fraction of patients with mutations in significant genes in the integrative topological analysis of 12 tumor types.** For each cancer type, the fraction of patients with a mutation in significant and significant actionable cancer associated genes is presented. Actionable genes were based on the list of actionable genes with approved drug of Bertrand *et al*.

| Cohort | Cancer-associated | Level 1 actionable |
|---|---|---|
| **BLCA** | 70% | 14% |
| **BRCA** | 76% | 0% |
| **CESC** | 23% | 0% |
| **COAD** | 100% | 88% |
| **GBM** | 47% | 39% |
| **HNSC** | 87% | 6% |
| **LGG** | 94% | 14% |
| **LUAD** | 88% | 14% |
| **PCPG** | 33% | 22% |
| **STAD** | 78% | 0% |
| **TGCT** | 34% | 17% |
| **THCA** | 75% | 63% |

**Supplementary Table 4 | TCGA data sources and parameters used in the analysis.** For each cohort, the version of the RSEM, MAF and MutSig2CV files used in the analysis is specified, as well as the threshold in the frequency of mutations that was used and the number of mutations (in $log_{10}$ scale) above which mutations where down-sampled (see Methods).

| Cohort | RSEM file | MAF file | Frequency threshold | Down-sampling | MutSig2CV (doi) |
|---|---|---|---|---|---|
| **BLCA** | 3.1.18.0 | Raw_Level_3.2015082100 | 0.060 | NA | 10.7908/C1MW2GGF |
| **BRCA** | 3.1.11.0 | Level_3.2015110100 | 0.015 | 2.3 | 10.7908/C1TB167Z |
| **CESC** | 3.1.11.0 | Level_3.2015110100 | 0.035 | 2.8 | 10.7908/C1MG7NV6 |
| **COAD** | 3.1.12.0 | Raw_Level_3.2015082100 | 0.060 | 3.0 | 10.7908/C1DF6QJD |
| **GBM** | 3.1.2.0 | Raw_Level_3.2015082100 | 0.040 | NA | 10.7908/C1XG9QGN |
| **HNSC** | 3.1.9.0 | Raw_Level_3.2015082100 | 0.040 | NA | 10.7908/C18C9VM5 |
| **LGG** | 3.1.13.0 | Raw_Level_3.2015082100 | 0.020 | 2.5 | 10.7908/C1MC8ZDF |
| **LUAD** | 3.1.14 | Raw_Level_3.2015082100 | 0.060 | NA | 10.7908/C17P8XT3 |
| **PCPG** | 3.1.2.0 | Level_3.2015110100 | 0.020 | NA | 10.7908/C13T9GN0 |
| **STAD** | 3.1.0.0 | Level_3.2015110100 | 0.040 | 2.6 | 10.7908/C1C828SM |
| **TGCT** | 3.1.0.0 | Level_3.2015110100 | 0.030 | NA | 10.7908/C1S1820D |
| **THCA** | 3.1.12.0 | Level_3.2015110100 | 0.009 | NA | 10.7908/C16W99KN |
